# YTHDC2 serves a distinct late role in spermatocytes during germ cell differentiation

**DOI:** 10.1101/2023.01.23.525146

**Authors:** Alexis S. Bailey, Margaret T. Fuller

**Author notes:** Corresponding author, lead contact for the manuscript: Margaret Fuller Department of Developmental Biology Beckman Center B300 279 Campus Drive Stanford University School of Medicine Stanford, CA 94305-5329. **Competing Interest:** The authors have no competing interests. **Classifications:** Biological Sciences / Developmental Biology.

## Abstract

Post-transcriptional regulation of gene expression by RNA-binding proteins can enhance the speed and robustness of cell state transitions by controlling RNA stability, localization, or if, when or where mRNAs are translated. The RNA helicase YTHDC2 is required to shut down components of the mitotic program to facilitate a proper switch from mitosis to meiosis in mouse germ cells. Here we show that YTHDC2 has a second essential role in promoting meiotic progression in late spermatocytes. Inducing conditional knockout of *Ythdc2* during the first wave of spermatogenesis, after initiation of meiotic prophase, allowed *Ythdc2*-deficient germ cells to advance to the pachytene stage and properly express many meiotic markers. However, the *Ythdc2*-deficient spermatocytes mis-expressed a number of genes, some up-regulated and some down-regulated, failed to transition to the diplotene stage, then quickly died. Co-immunoprecipitation experiments revealed that YTHDC2 interacts with several RNA-binding proteins in early or late spermatocytes, with many of the interacting proteins, including MEIOC, localizing to granules, similar to YTHDC2. Our findings suggest that YTHDC2 collaborates with other RNA granule components to facilitate proper progression of germ cells through multiple steps of meiosis via mechanisms influencing post-transcriptional regulation of RNAs.

**SIGNIFICANCE STATEMENT:** An effective, robust switch from mitosis to meiosis is essential for the production of gametes in sexually reproducing organisms. The RNA helicase YTHDC2 is required for germ cells to shut down aspects of the mitotic program as they initiate meiotic prophase in the mouse male germline. Here we utilize a timed conditional knockout strategy to show that, in addition to its conserved function in the mitosis-to-meiosis transition, YTHDC2 has a second critical role in promoting the pachytene-to-diplotene transition late in male meiotic prophase. YTHDC2 interacts with several proteins that are also present in RNA granules, including MEIOC, suggesting that YTHDC2 collaborates with RNA granule components to regulate RNAs as germ cells progress from one cell state to the next.

## INTRODUCTION

RNA-binding proteins and post-transcriptional regulation of RNAs play fundamental roles in germ cell development, allowing an efficient transition from the mitotic to the meiotic program and progression of germ cells through meiotic prophase and terminal differentiation. The DExH-box RNA helicase YTHDC2 is the mammalian homolog of *Drosophila* Benign gonial cell neoplasm (Bgcn), which along with its protein binding partners Bag of marbles (Bam) and Tumorous testis (Tut), regulates the switch from mitotic divisions to meiosis in *Drosophila* males. Both Bgcn and Tut have RNA-binding domains, and the complex has been shown to interact with at least one target mRNA through binding its 3′ UTR (1, 2). Previous studies from our lab and others have shown that YTHDC2 function is required to properly execute the transition from mitosis to meiosis in the mouse germline (3–7). *Ythdc2* null mutant male germ cells enter meiotic prophase and start to express some meiotic factors, but fail to turn down expression of certain mitotic regulators. The resulting null mutant early spermatocytes have a mixed identity, prematurely condense their chromosomes in a mitosis-like metaphase array, and then quickly die by apoptosis (3, 8). As YTHDC2 localizes to the cytoplasm and has been shown to directly bind specific transcripts, including the mitotic cyclin *Ccna2* (3, 9), YTHDC2 likely regulates gene expression post-transcriptionally, similar to *Drosophila Bgcn*.

Previous studies have shown that YTHDC2 binds the meiotic protein MEIOC (3, 8, 10, 11) as well as the RNA-binding protein RBM46 (8, 12). Binding of YTHDC2 to both MEIOC and RBM46 did not require RNA. *Meioc* and *Rbm46* knockout males showed the same distinctive early spermatocyte death phenotype as *Ythdc2* null mutants (10–13), suggesting that YTHDC2, MEIOC and RBM46 function together to regulate the switch from mitosis to meiosis in males. Directly tying YTHDC2 target RNAs to degradation machinery and supporting a functional role for YTHDC2 in regulating RNA stability, previous studies also showed that the cytoplasmic 5′ → 3′ exoribonuclease XRN1 is a direct, RNA-independent binding partner of YTHDC2 (7, 8, 14).

YTHDC2 protein expression is strongly up-regulated as male germ cells start meiosis. Analysis by western blot indicated that YTHDC2 protein levels continue to increase during the first wave of spermatogenesis as germ cells proceed through meiotic prophase (3). As *Ythdc2* null mutant germ cells die very soon after entry into meiotic prophase, it was not previously possible to investigate whether YTHDC2 has functional roles in male germ cell differentiation after its role in shutting down the mitotic program. It was also unclear whether the abnormally low levels of many meiotic RNAs such as *Spo11* observed in testes from *Ythdc2* null mutant males could be a secondary effect of the death of young spermatocytes early in meiotic prophase. Here we establish conditions that allow YTHDC2 to act in very early spermatocytes, assuring that the cells escape the attempted mitotic division and early apoptosis characteristic of null mutants, but then knock out expression by the pachytene spermatocyte stage to test the role of YTHDC2 in later stages of germ cell differentiation.

Our results show that in addition to its role in the mitosis-to-meiosis transition, YTHDC2 also has a critical function in promoting progression of spermatocytes through late meiotic prophase. To carefully examine the direct consequences of *Ythdc2* loss of function in late spermatocytes, we induced conditional knockout of *Ythdc2* once the first wave of germ cells reached the early spermatocyte stage. This strategy allowed us to largely bypass the mitosis-to-meiosis defects previously observed in *Ythdc2* null mutants and examine the requirement for YTHDC2 function at later stages of meiotic prophase, when expression of the protein normally peaks. Induced deletion of *Ythdc2* in early spermatocytes allowed the germ cells to properly progress to the pachytene stage, form synaptonemal complexes and robustly express meiotic markers. However, loss of function of *Ythdc2* in spermatocytes resulted in changes in expression level for a number of genes, some up-regulated and some down-regulated, followed by elimination of late pachytene spermatocytes by apoptosis prior to the meiotic divisions.

Immunoprecipitation (IP)-mass spectrometry experiments from first wave testes containing early or late spermatocytes indicated that YTHDC2 interacts with several other RNA-binding proteins that play roles in RNA metabolism. Follow-up co-IP-western blots revealed that some of the protein interactions were RNA-independent while others depended on the presence of RNA. Many of the interacting proteins have been shown to localize to perinuclear granules, similar to YTHDC2. Strikingly, we find that MEIOC localizes to granules in pachytene spermatocytes, and that the localization of MEIOC to granules was enhanced in the *Ythdc2* cKO. Taken together, our results indicate that YTHDC2 functions together with other RNA granule components through post-transcriptional regulation of RNAs to facilitate proper progression of the meiotic program.

## RESULTS

### YTHDC2 is highly expressed in late spermatocytes

Immunofluorescence staining detected YTHDC2 protein starting in early spermatocytes, with expression continuing throughout meiotic prophase, as previously reported (3, 5). Similar immunofluorescence staining of cross-sections of testis tubules from males 18 days after birth (P18) showed low levels of YTHDC2 signal in the cytoplasm of early leptotene spermatocytes (second wave) (Fig. 1A, white arrowheads) and high levels of cytoplasmic YTHDC2 signal in the late pachytene spermatocytes (first wave) that occupy the center of the testis tubules (Fig. 1A, white arrows), indicating that YTHDC2 protein expression increases as spermatocytes mature. This raised the question of whether YTHDC2 has a critical function in late spermatocytes in addition to its role in early meiotic prophase.

**Fig. 1.**
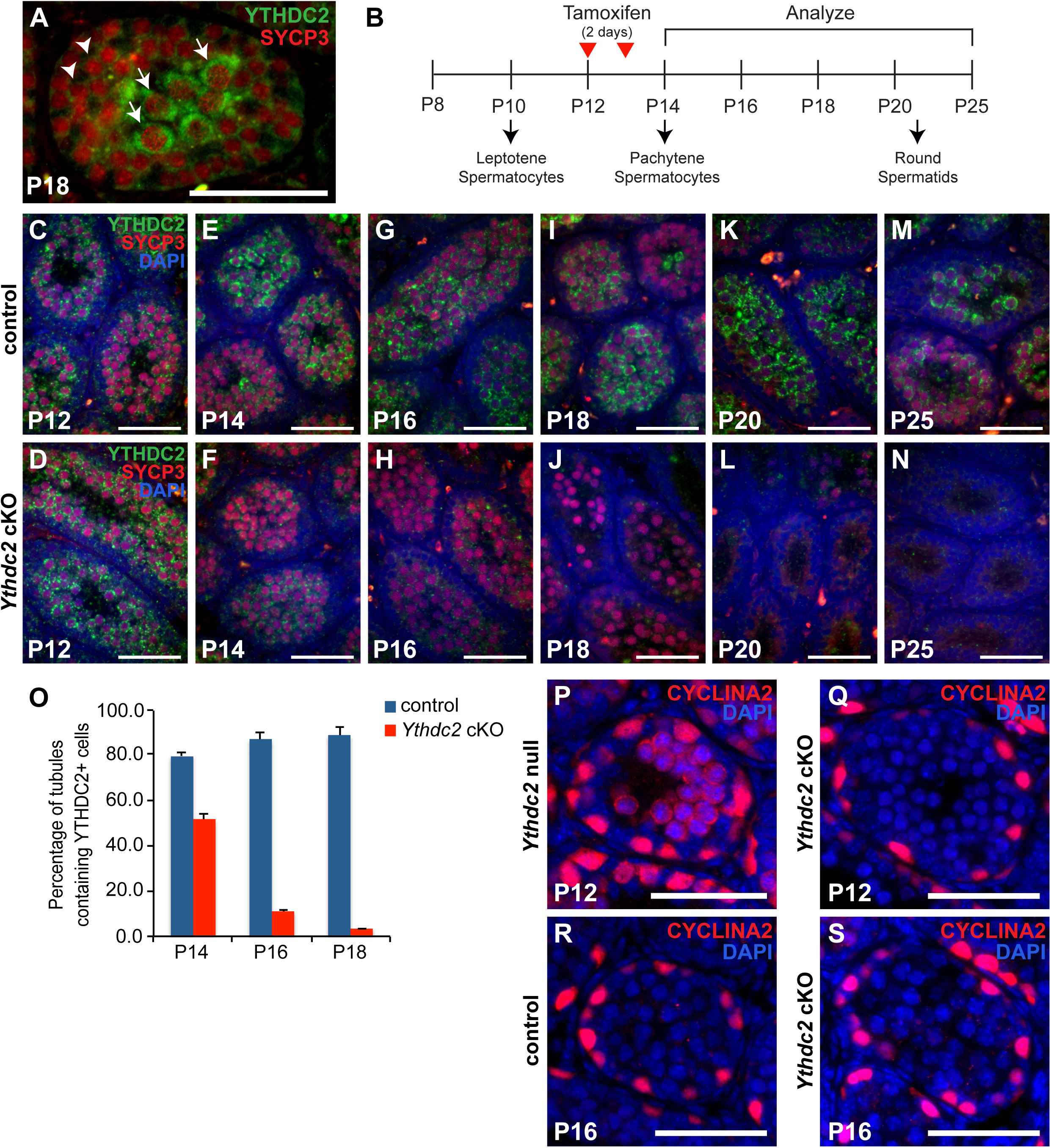
Knock out of *Ythdc2* in spermatocytes during the first wave of spermatogenesis. (A) Immunofluorescence image of a P18 wild-type mouse testis tubule cross-section stained for YTHDC2 (green) and SYCP3 (red). Arrowheads: leptotene spermatocytes from the second wave of spermatogenesis. Arrows: pachytene spermatocytes from the first wave of spermatogenesis within the lumen of a testis tubule. (B) Timeline of initial appearance of germ cell stages during the first wave of spermatogenesis, with the timing of tamoxifen injections given to both control and *Ythdc2* cKO mice. (C-N) Immunofluorescence images of control (*+/+; Ythdc2^flox/Δ^*, top) and *Ythdc2* cKO (*UBC-CreERT2; Ythdc2^flox/Δ^*, bottom) seminiferous tubule cross-sections from (C and D) P12, (E and F) P14, (G and H) P16, (I and J) P18, (K and L) P20 and (M and N) P25 testes stained with anti-YTHDC2 (green) and anti-SYCP3 (red) antibodies as well as DAPI to mark the DNA (blue). (O) Percentage of tubules containing YTHDC2-positive cells in P14, P16 and P18 control (+/+*; Ythdc2^flox/Δ^*) (blue) and *Ythdc2* cKO (red) testes (n = 2-4 mice per group; *t* test, P14: p-value = 0.0003, P16: p-value < 0.0001, P18: p-value = 0.0019). Error bars: SEM. (P-S) Immunofluorescence images of testis tubules from (P) P12 *Ythdc2* null, (Q) P12 *Ythdc2* cKO, (R) P16 control (*+/+; Ythdc2^flox/Δ^*) and (S) P16 *Ythdc2* cKO mice stained with anti-CYCLINA2 (red) antibody and DAPI (blue). Scale bars: 50 µm.

To bypass the early *Ythdc2* null mutant phenotype and examine the function of YTHDC2 in late meiotic prophase, we generated mice bearing a conditional *Ythdc2* allele (*Ythdc2^flox^*) (Fig. S1) with *loxP* sites flanking exons 6 and 7. *Ythdc2^flox^* mice were crossed to our previously generated mice carrying the *Ythdc2* null allele (*Ythdc2^Δ^*) (3). We then introduced the tamoxifen-inducible transgene *UBC-Cre-ERT2* (15), which expresses CRE-ERT2 under the control of the *ubiquitin C* (*UBC*) promoter. Treatment of *UBC-Cre-ERT2; Ythdc2^flox/Δ^* (*Ythdc2* cKO) juvenile mice with tamoxifen at P12 and P13, when the first wave of germ cells have already entered meiotic prophase and are in the leptotene/zygotene stage (Fig. 1B), allowed expression of YTHDC2 protein in early-stage spermatocytes but substantially reduced levels of YTHDC2 protein in late-stage spermatocytes. YTHDC2 protein was detected at similar levels by immunofluorescence staining in control (no *Cre*) and *Ythdc2* cKO early spermatocytes at P12, the time of the first tamoxifen injection (Fig. 1C and D). YTHDC2 protein levels started to decrease one day after the second tamoxifen injection (P14) in seminiferous tubules from *Ythdc2* cKO males compared to control males (Fig. 1F vs. E), although low levels of YTHDC2 protein were still detected in approximately 50% of the *Ythdc2* cKO tubules (Fig. 1O). By P16, while spermatocytes from tamoxifen-treated control males lacking the *Cre* transgene continued to have high levels of YTHDC2 protein expression (Fig. 1G), the level of YTHDC2 protein in spermatocytes in *Ythdc2* cKO testis tubules detected by immunofluorescence staining was considerably lower (Fig. 1H). Quantification of the percentage of tubules that contained YTHDC2-positive spermatocytes confirmed that YTHDC2 protein expression was below the level of detection by our immunofluorescence staining in the majority of *Ythdc2* cKO testis tubules by P16 (Fig. 1O).

### YTHDC2 is required for late spermatocytes to progress to post-pachytene stages

Analysis of testis tubule cross-sections from *Ythdc2* cKO males showed that the timing of tamoxifen treatment we employed allowed male germ cells to proceed past the initial requirement for YTHDC2 function early in meiotic prophase. Our previous studies revealed that in *Ythdc2* null mutant mice, a large number of germ cells underwent abnormal chromosome condensation soon after initiating meiotic prophase (3), visible as abnormal metaphase-like spermatocytes in P12 testes. This phenotype was also observed in *Meioc* and *Rbm46* knockout males (10–12). Quantification of testis tubule cross-sections revealed that only a small percentage of tubules in *Ythdc2* cKO mice contained cells with condensed chromosomes at either P16 (no-*Cre* control 1.4% vs. *Ythdc2* cKO 8.7%) or P18 (no-*Cre* control 0% vs. *Ythdc2* cKO 3.9%).

Consistent with the small number of cells with abnormal chromosomal condensation in *Ythdc2* cKO mice, *Ythdc2* cKO spermatocytes did not express the abnormally high levels of CYCLINA2 previously observed in P12 *Ythdc2* null mutant spermatocytes (Fig. 1Q and S vs. P) (3), indicating that inducing knockout of *Ythdc2* starting at P12 allowed the majority of germ cells to get past the requirement for YTHDC2 to shut down the mitotic program during the first wave of spermatogenesis.

Examination of histological sections of P16 testes showed that, unlike *Ythdc2* null mutant germ cells, which never reached the pachytene spermatocyte stage (3), *Ythdc2* cKO germ cells moved successfully through the early spermatocyte stages and reached pachytene at a similar frequency to control mice lacking the *Cre* transgene but treated with the same tamoxifen regimen (Fig. 2E, F and Q). Immunofluorescence staining revealed that P16 *Ythdc2* cKO tubules were filled with pachytene spermatocytes that expressed high levels of the synaptonemal complex component SYCP3 and had nuclear puncta of γH2AX (Fig. 2N and P) although they lacked detectable YTHDC2 immunofluorescence signal.

**Fig. 2.**
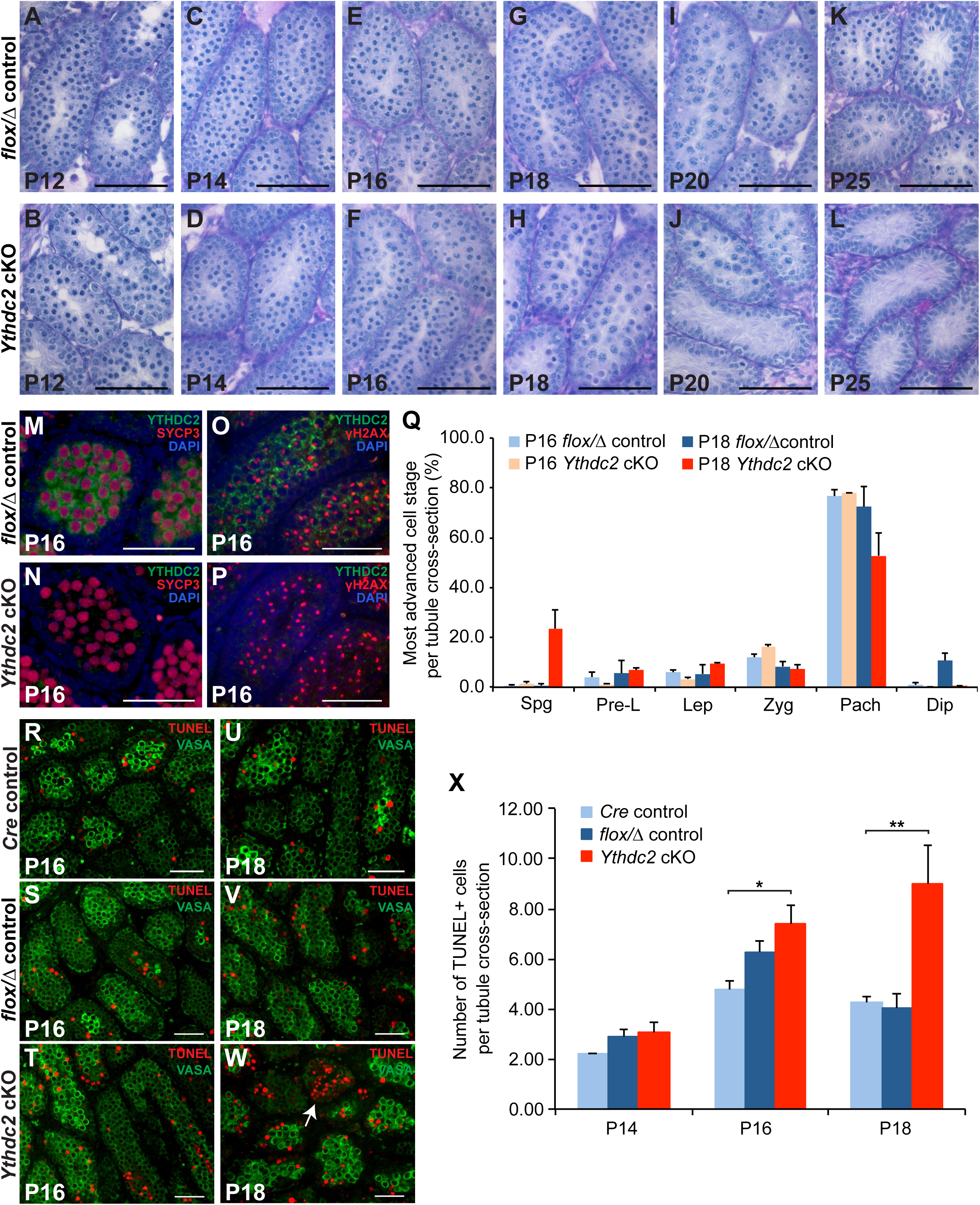
YTHDC2 is required for late spermatocytes to progress through meiotic prophase. (A-L) Sections of mouse seminiferous tubules stained with periodic acid-Schiff (PAS) from (A and B) P12, (C and D) P14, (E and F) P16, (G and H) P18, (I and J) P20 and (K and L) P25 tamoxifen-treated control (*+/+; Ythdc2^flox/Δ^*, top) and *Ythdc2* cKO (*UBC-CreERT2; Ythdc2^flox/Δ^*, bottom) testis. Scale bars: 50 µm. (M-P) Testis from P16 control (*+/+; Ythdc2^flox/Δ^*) and *Ythdc2* cKO (*UBC-CreERT2; Ythdc2^flox/Δ^*) mice stained with (M and N) anti-YTHDC2 (green), anti-SYCP3 (red) antibodies as well as DAPI to mark the DNA (blue) or (O and P) anti-YTHDC2 (green), anti-γH2AX (red) antibodies and DAPI (blue). (Q) Percentage of tubule cross-sections containing spermatogonia (Spg), preleptotene (Pre-L), leptotene (Lep), zygotene (Zyg), pachytene (Pach) or diplotene (Dip) as the most advanced cell stage in P16 and P18 control (+/+*; Ythdc2^flox/Δ^*) and *Ythdc2* cKO (*UBC-CreERT2; Ythdc2^flox/Δ^*) testes (n = 2 mice per group; > 120 tubules counted per mouse). Error bars: SEM. (R-W) Testis tubule cross-sections from (R-T) P16 and (U-W) P18 *Cre* control (*UBC-CreERT2; Ythdc2^flox/+^*), *flox/Δ* control (+/+*; Ythdc2^flox/Δ^*), and *Ythdc2* cKO (*UBC-CreERT2; Ythdc2^flox/Δ^*) mice labeled with TUNEL (red) and VASA (green). Arrow: Tubule filled with TUNEL-positive cells. Scale bars: 50 µm. (X) Number of TUNEL-positive cells per tubule cross-section in P14, P16 and P18 *Cre* control (light blue), *flox/Δ* control (dark blue), and *Ythdc2* cKO (red) testes (n = 2-4 mice per group; > 110 tubules counted per mouse; *t* test, * p-value = 0.0176, ** p-value = 0.0268). Error bars: SEM.

While *Ythdc2* cKO pachytene spermatocytes appeared normal in histological preparations, the cells failed to advance past the late pachytene stage and quickly underwent apoptosis, suggesting that YTHDC2 is required for late spermatocytes to complete meiotic prophase. Histological analysis of tubule cross-sections showed that while some tubules from tamoxifen-treated control males contained diplotene spermatocytes by P18 (Fig. 2Q), *Ythdc2* cKO germ cells progressed to the diplotene stage very rarely (one diplotene cell in > 500 tubule cross-sections). Rather, primarily late pachytene spermatocytes were present in the P18 *Ythdc2* cKO tubules (Fig. 2H and Q). By P20, while control tubules were filled with late-stage spermatocytes (Fig. 2I), the majority of tubules in *Ythdc2* cKO mice lacked late spermatocytes (Fig. 2J). *Ythdc2* cKO tubules were empty of all spermatocyte stages by P25 (Fig. 2L). TUNEL staining showed a slight increase in the number of TUNEL-positive cells starting at P16 in *Ythdc2* cKO testes (Fig. 2T) compared to controls lacking either the *Ythdc2* null allele (*UBC-Cre-ERT2; Ythdc2^flox/+^*; *Cre* control) or the *Cre* transgene (*+/+; Ythdc2^flox/Δ^* ; *flox/Δ* control) treated with the same tamoxifen regimen (Fig. 2R and S, quantified in X). However, the number of TUNEL-positive cells per tubule cross-section greatly increased at P18 in *Ythdc2* cKO testes, with some *Ythdc2* cKO tubules filled with TUNEL-positive spermatocytes (Fig. 2W, white arrow), compared to either *Cre* control or *flox/Δ* control testes (Fig. 2U and V, quantified in X). The number of TUNEL-positive cells per tubule cross-section was similarly low/comparable in both negative controls: (1) mice carrying the *Cre* transgene but not the *Ythdc2* null allele and (2) mice carrying *Ythdc2^flox/Δ^* but lacking the *Cre* transgene, indicating that the death of late spermatocytes prior to diplotene was not due to *Cre* induced toxicity.

Detailed analysis by immunofluorescence staining for several meiotic markers confirmed that *Ythdc2* cKO germ cells progressed to the pachytene stage. SYCP3 properly localized on the chromosome axis in early (Fig. 3B and D) and late (Fig. 3F and H) *Ythdc2* cKO spermatocytes, similar to spermatocytes from tamoxifen-treated control males lacking the *Cre* transgene. *Ythdc2* cKO early spermatocytes appeared to properly make double strand breaks, as immunofluorescence staining of chromosome spreads showed γH2AX foci along the chromosomes in leptotene (Fig. 3B) and zygotene (Fig. 3D) spermatocytes. In pachytene spermatocytes, γH2AX became restricted to the sex vesicle containing the X and Y chromosomes in *Ythdc2* cKO males (Fig. 3F), similar to no-*Cre* control males (Fig. 3E).

**Fig. 3.**
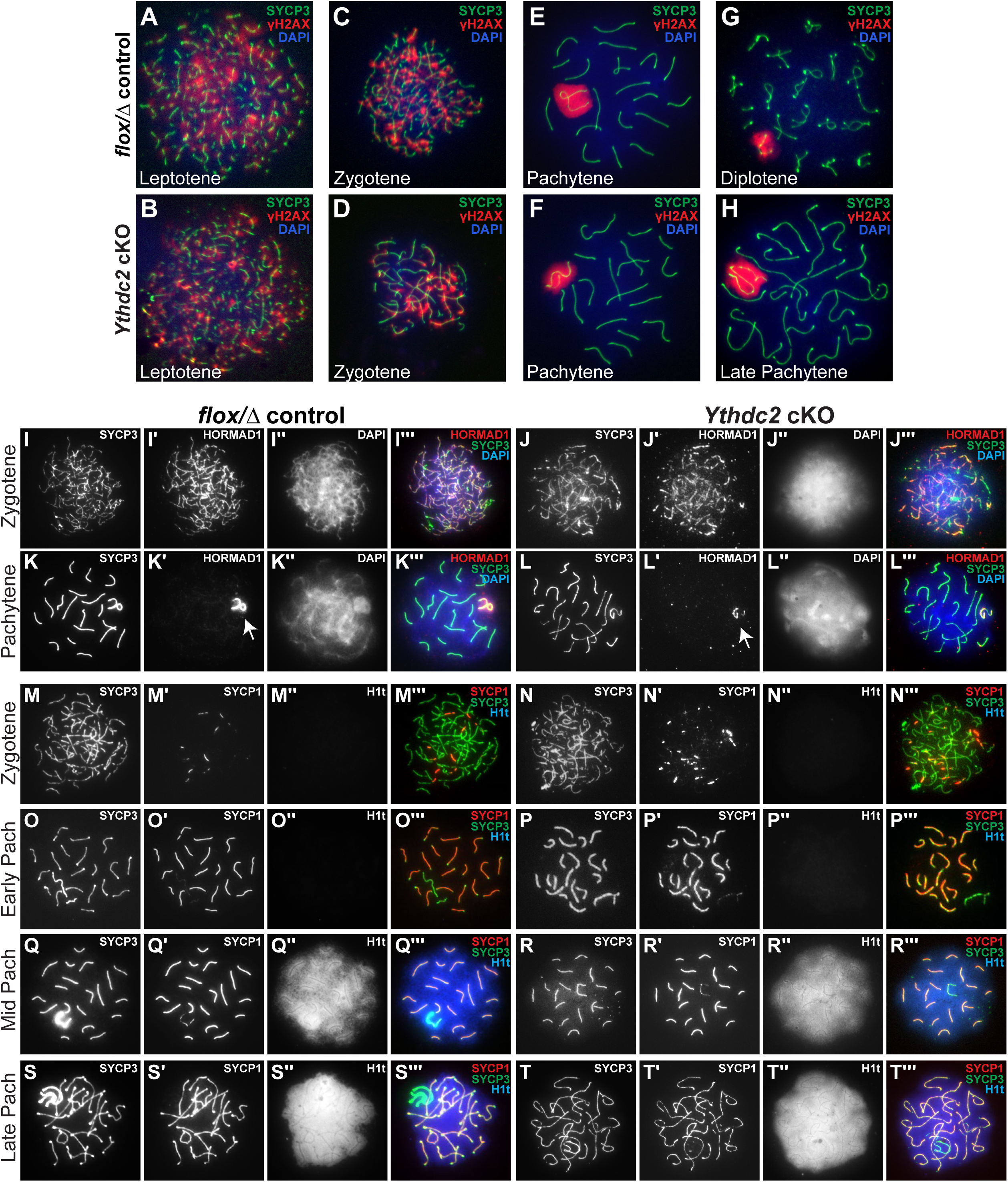
Spermatocytes lacking YTHDC2 reach the late pachytene stage but then quickly die. (A-H) Immunofluorescence images of germ cell spreads from P18 tamoxifen-treated control (*+/+; Ythdc2^flox/Δ^*, top) and *Ythdc2* cKO (*UBC-CreERT2; Ythdc2^flox/Δ^*, bottom) mice stained with anti-SYCP3 (green), anti-γH2AX (red) antibodies and DAPI to mark the DNA (blue). (I-Lʹʹʹ) Immunofluorescence images of germ cell spreads from P16 control (*+/+; Ythdc2^flox/Δ^*, left) and *Ythdc2* cKO (*UBC-CreERT2; Ythdc2^flox/Δ^*, right) mice stained with anti-SYCP3 (green), anti-HORMAD1 (red) antibodies and DAPI (blue). Arrows: HORMAD1 staining on the sex body in pachytene spermatocytes. (M-Tʹʹʹ) Germ cell spreads from P16 control (*+/+; Ythdc2^flox/Δ^*, left) and *Ythdc2* cKO (*UBC-CreERT2; Ythdc2^flox/Δ^*, right) mice stained with anti-SYCP3 (green), anti-SYCP1 (red) and anti-H1t (blue) antibodies.

Further characterization of pachytene spermatocyte substages in germ cell spreads revealed that early, mid and late pachytene spermatocytes were present in *Ythdc2* cKO testes and that the temporal ordering of key meiotic events was executed correctly. HORMAD1 co-localized with SYCP3 in P16 *Ythdc2* cKO zygotene spermatocytes (Fig. 3J-Jʹʹʹ) as in no-*Cre* controls (Fig. 3I-Iʹʹʹ), then was properly depleted from the chromosome axes after SC formation, with expression remaining on the unsynapsed regions of the sex chromosomes in *Ythdc2* cKO pachytene spermatocytes, similar to control males (Fig. 3K-Lʹʹʹ, white arrows). The transverse filament protein SYCP1 started to localize at synapsing homologous chromosomes in *Ythdc2* cKO zygotene spermatocytes (Fig. 3N-Nʹʹʹ) as in controls (Fig. 3M-Mʹʹʹ). The major linker histone variant in pachytene spermatocytes, H1t, was absent in early pachytene spermatocytes (Fig. 3P-Pʹʹʹ), started to be present in mid-pachytene spermatocyte nuclei (Fig. 3R-Rʹʹʹ) and was abundant in late pachytene spermatocytes (Fig. 3T-Tʹʹʹ) in the *Ythdc2* cKO, as in no-*Cre* control spermatocytes (Fig. 3O, Q and S). Analysis of the dynamics of RAD51 expression revealed that RAD51 foci were present at the zygotene stage (Fig. S2A and B) then decreased in number in *Ythdc2* cKO pachytene spermatocytes (Fig. S2D and F), similar to control males (Fig. S2C and E), suggesting that the double strand breaks were being properly repaired.

In addition to the inducible *Ythdc2* knockout model, *Ythdc2^flox/Δ^* conditional mice were also crossed to mice carrying a *Spo11-Cre* transgene, which expresses CRE recombinase in spermatocytes that have initiated meiosis (16). Consistent with the findings from the inducible *UBC-Cre-ERT2; Ythdc2^flox/Δ^*mice, examination of P18 control (*Spo11-Cre*; *Ythdc2^flox/+^*) and *Spo11-Cre; Ythdc2^flox/Δ^* testis tubules revealed that YTHDC2 is required to complete meiotic prophase. *Spo11-Cre; Ythdc2^flox/Δ^* germ cells reached the pachytene stage of meiotic prophase, as scored by immunofluorescence staining for γH2AX, which was restricted to the sex body as in control pachytene spermatocytes (Fig. S3C and D). However, the spermatocytes then quickly started to die, leading to some tubules lacking late spermatocytes in P18 *Spo11-Cre; Ythdc2^flox/Δ^* testes (Fig. S3B). Furthermore, analysis of histological sections from adult testes revealed that while the *Spo11-Cre; Ythdc2^flox/Δ^* model was not as penetrant as the inducible system, a large proportion of tubules lacked late-stage germ cells (Fig. S3F vs. E), and mature sperm were rarely seen in the epididymis (Fig. S3H vs. G). Taken together, our data from analysis of both the cell-stage specific and the inducible *Ythdc2* conditional knockout mice revealed that in addition to its early role at the mitotic-to-meiotic transition, YTHDC2 has a distinct, later role in meiotic prophase required for pachytene spermatocytes to successfully become diplotene spermatocytes.

### YTHDC2 interacts with several other proteins involved in RNA regulation

Consistent with a role in RNA regulation, YTHDC2 appears to interact with several proteins involved in RNA biology, including RNA stability, RNA localization, and translation.

Immunoprecipitation of YTHDC2 from P12 or P18 wild-type testes followed by mass spectrometry revealed several candidate interacting protein partners for YTHDC2 (Fig. S4A and B). Co-immunoprecipitation in the presence or absence of RNase validated candidate interacting proteins and revealed which could interact with YTHDC2 independently of RNA. Multiple proteins co-immunoprecipitated with YTHDC2 from RNase-treated testis extracts, including the previously known interacting protein MEIOC, as well as several additional proteins, including PABPC1 and FMR1 (Fig. 4A). MEIOC, PABPC1 and FMR1 showed particularly strong binding to YTHDC2 independent of RNA. At P18, CAPRIN1 and EWS also co-immunoprecipitated with YTHDC2 in the presence of RNase, although much more was recovered with YTHDC2 when RNA was present (Fig. 4A). The hnRNP PTBP1 appeared to co-immunoprecipitate with YTHDC2 at low levels in the presence of RNase at P12 but was not detected when YTHDC2 was immunoprecipitated in the presence of RNase from P18 testes. Four additional proteins co-immunoprecipitated with YTHDC2, but only in the presence of RNA: MOV10, YBX1, G3BP2 and UPF1 (Fig. 4B). Immunofluorescence staining of YTHDC2 and the protein-binding partners in wild-type testes showed that the interactors were all highly expressed in the cytoplasm at specific spermatocyte stages (Fig. S4C-L). Some of the interacting proteins, including PTBP1, MOV10 and G3BP2, were expressed at high levels primarily in early leptotene and zygotene spermatocytes, while others, including PABPC1, CAPRIN1, YBX1 and UPF1, were more highly expressed in pachytene spermatocytes, suggesting that YTHDC2 could associate with different protein complexes at different stages.

**Fig. 4.**
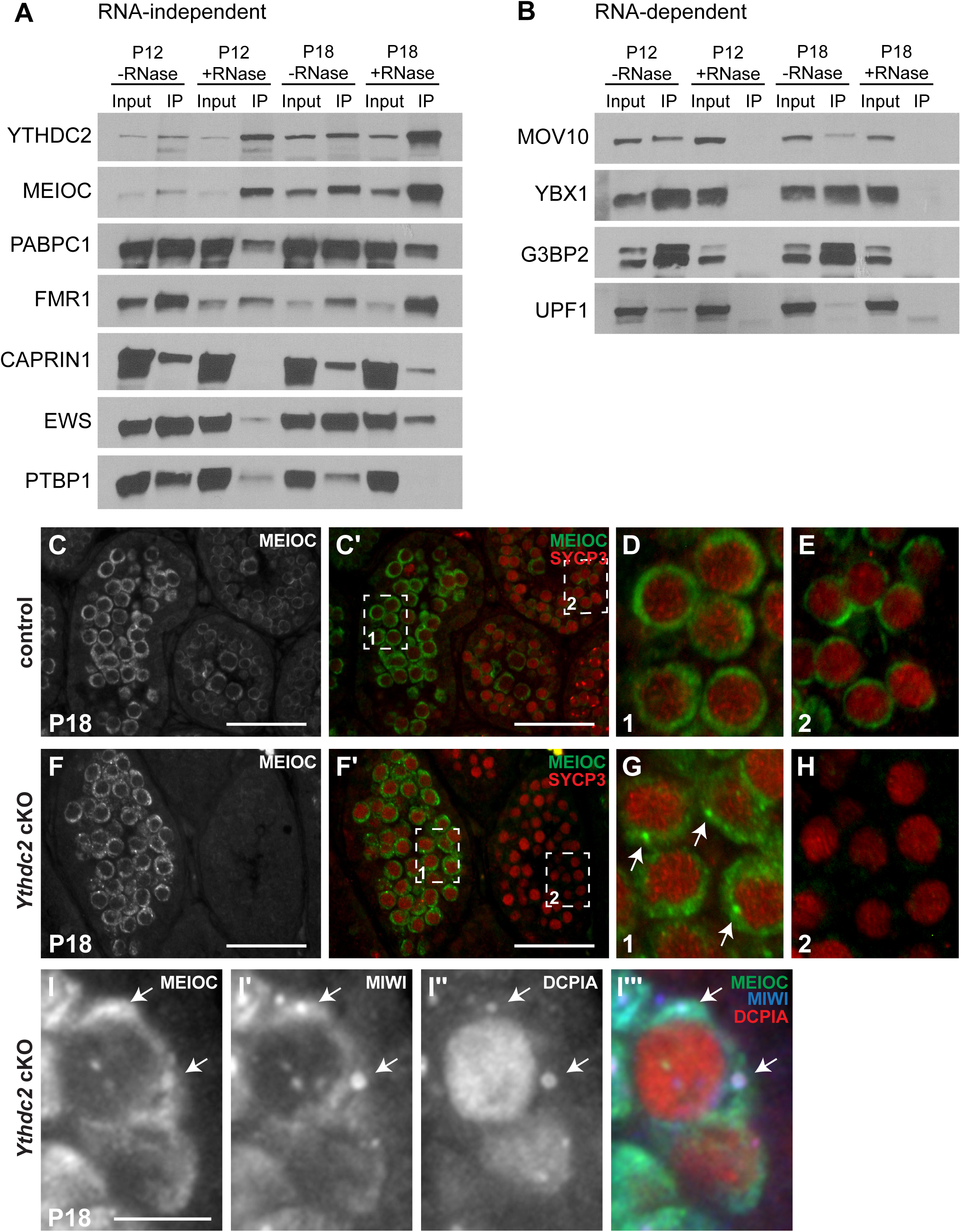
YTHDC2 interacts with several other proteins involved in RNA regulation. (A and B) Immunoprecipitation with α-YTHDC2 from P12 and P18 wild-type mouse testis extracts treated either with RNase Inhibitor (-RNase) or RNase A (+RNase). Western blots probed with anti-YTHDC2, anti-MEIOC, anti-PABPC1, anti-FMR1, anti-CAPRIN1, anti-EWS, anti-PTBP1, anti-MOV10, anti-YBX1, anti-G3BP2 and anti-UPF1 antibodies. (C-H) Immunofluorescence images of testis cross-sections from P18 (C-E) control (*+/+; Ythdc2^flox/Δ^*) and (F-H) *Ythdc2* cKO (*UBC-CreERT2; Ythdc2^flox/Δ^*) mice stained with anti-MEIOC (green) and anti-SYCP3 (red) antibodies. Scale bars: 50 µm. (D and E) High-magnification images of (D) pachytene spermatocytes (first wave) in boxed region 1 in panel C′ and (E) leptotene spermatocytes (second wave) in boxed region 2 in panel C′. (G and H) High-magnification images of (G) pachytene spermatocytes (first wave) in boxed region 1 in panel F′ and (H) leptotene spermatocytes (second wave) in boxed region 2 in panel F′. Arrows: MEIOC protein in perinuclear puncta in pachytene spermatocytes. (I-Iʹʹʹ) High-magnification immunofluorescence images of spermatocytes from a P18 *Ythdc2* cKO mouse testis cross-section stained with anti-MEIOC (green), anti-MIWI (blue) and anti-DCP1A (red) antibodies. Arrows: puncta in the cytoplasm of pachytene spermatocytes. Scale bar: 5 µm.

MEIOC protein strongly co-immunoprecipitated with YTHDC2 from both P12 and P18 testes in the absence of RNA. Immunofluorescence staining of control versus *Ythdc2* cKO P18 testes with anti-MEIOC and anti-SYCP3 revealed that loss of YTHDC2 affected the subcellular distribution of MEIOC protein. P18 testis tubules contained both pachytene spermatocytes from the first wave of spermatogenesis (Fig. 4C′ and F′, box 1) and leptotene spermatocytes from the second wave (Fig. 4C′ and F′, box 2). In control testes, MEIOC protein was detected in both pachytene (Fig. 4D) and leptotene (Fig. 4E) spermatocytes as previously shown (10, 11).

MEIOC protein was detected in P18 *Ythdc2* cKO pachytene spermatocytes (Fig. 4G). However, MEIOC protein was not detected by immunofluorescence in *Ythdc2* cKO leptotene spermatocytes from the second wave (Fig. 4H), consistent with our previous findings that MEIOC protein was expressed at much lower levels in leptotene spermatocytes in P12 *Ythdc2* null mutant testes compared to wild type (3). Taken together, these data suggest that YTHDC2 may promote the stability of MEIOC protein in early spermatocytes.

We had previously shown that YTHDC2 protein localizes to RNA granules in spermatocytes (3). While MEIOC protein was detected throughout the cytoplasm in *Ythdc2* cKO pachytene spermatocytes, it appeared strongly concentrated in perinuclear puncta (Fig. 4G, white arrows). In contrast, MEIOC protein did not appear to accumulate as strongly in perinuclear puncta in control P18 pachytene spermatocytes, but was more distributed throughout the cytoplasm (Fig. 4D). Co-staining with antibodies against two RNA granule components, the decapping enzyme DCP1A and the mouse PIWI family protein MIWI, revealed that many of the MEIOC-positive cytoplasmic puncta present in *Ythdc2* cKO spermatocytes also contained DCP1A and MIWI (Fig. 4I-Iʹʹʹ, white arrows).

### Early transcript changes following conditional knockout of *Ythdc2* in first wave spermatocytes

Comparison of RNAs expressed in *Ythdc2* cKO versus age-matched *Cre* control (*UBC-CreERT2; Ythdc2^flox/+^*) or *flox/Δ* control (+/+*; Ythdc2^flox/Δ^*) testes at P14 and P16 by RNA sequencing revealed early changes in gene expression before *Ythdc2* knockout spermatocytes began to die. In all cases, the mice were treated with two days of tamoxifen injections, at P12 and P13. While *Ythdc2* RNA expression was significantly down-regulated by P14 in the *Ythdc2* cKO testes compared to either the *Cre* (Fig. 5A) or *flox/Δ* control (Fig. S5B), expression of other genes in *Ythdc2* cKO testes were similar to both controls at P14 (Fig. 5A and Fig. S5A and B). At the P14 time point, only one day post the second tamoxifen injection, levels of YTHDC2 protein detected by immunofluorescence staining were decreasing in *Ythdc2* cKO testis tubules, but YTHDC2 protein was still detected in many tubules (Fig. 1F and O).

**Fig. 5.**
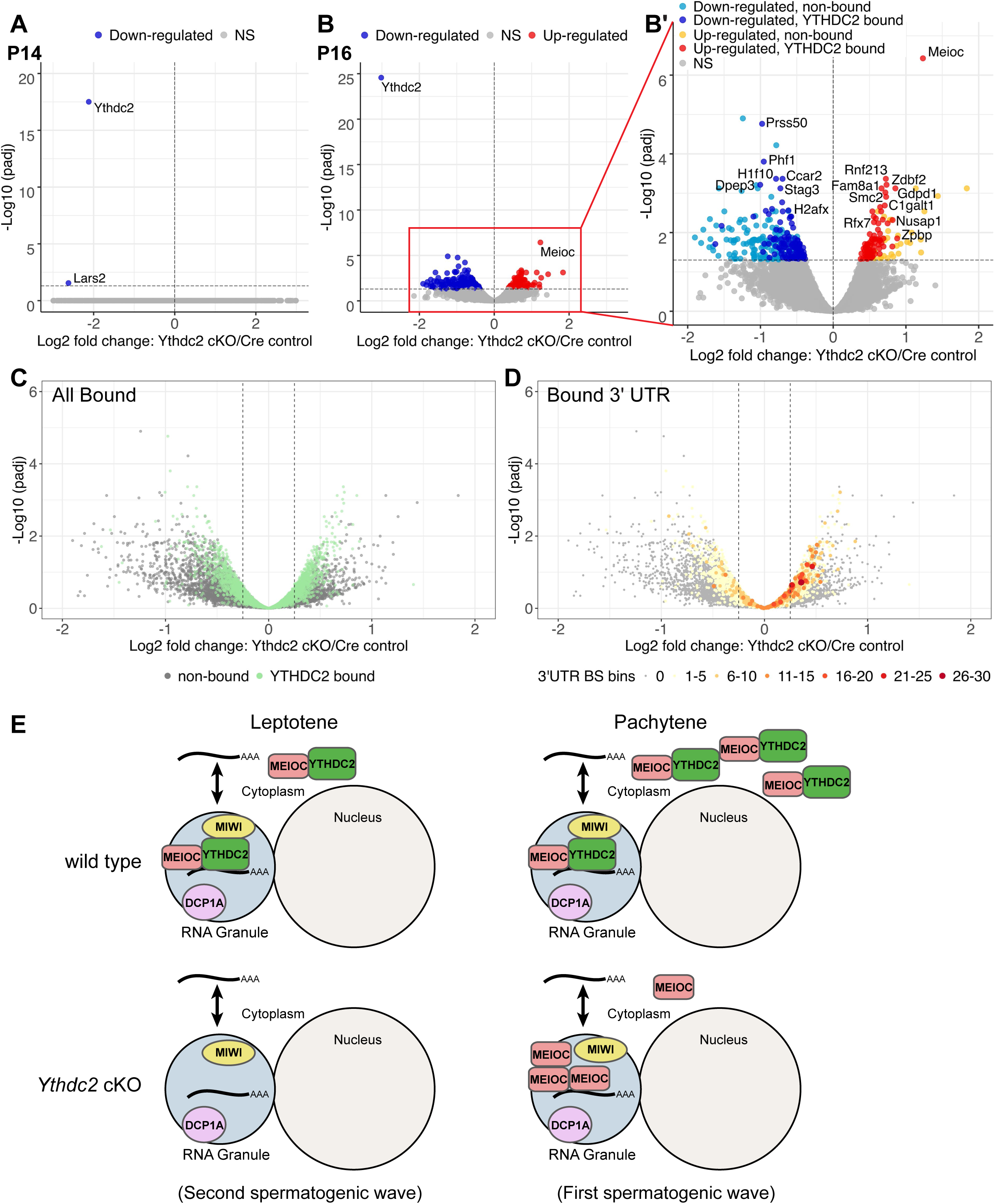
Early transcript changes following conditional knockout of *Ythdc2* in first wave spermatocytes. (A and B) Volcano plots representing significantly differentially expressed genes (adjusted p-value < 0.05, horizontal dashed line) in *Ythdc2* cKO (*UBC-CreERT2; Ythdc2^flox/Δ^*) testes relative to *Cre* control (*UBC-CreERT2; Ythdc2^flox/+^*) testes at (A) P14 and (B) P16. (Red) genes up-regulated, and (Blue) genes down-regulated in *Ythdc2* cKO testes relative to *Cre* control testes. (B′) Enlarged plot of region in the red box in panel B. Genes up and down-regulated in *Ythdc2* cKO testes color-coded based on whether the encoded transcripts were previously identified as bound by YTHDC2 based on CLIP (8). (Light blue) down-regulated, non-bound; (Dark blue) down-regulated YTHDC2 bound; (Yellow) up-regulated, non-bound; (Red) up-regulated, YTHDC2 bound. (C) Volcano plot depicting fold changes in gene expression in P16 *Ythdc2* cKO testes relative to *Cre* control testes for (green) all YTHDC2-bound RNAs identified by YTHDC2 CLIP (8) versus (gray) non-bound RNAs. (D) Volcano plot depicting fold changes in gene expression in *Ythdc2* cKO testes relative to *Cre* control testes. RNAs identified through CLIP as bound by YTHDC2 at their 3′ UTR by Li et al. (2022) are highlighted using a gradient color scale representing the number of YTHDC2 binding sites (BS) in the 3′ UTR. Dashed gray lines indicate Log2FC -0.25 and Log2FC 0.25. (E) Model depicting YTHDC2 and MEIOC protein expression and localization in RNA granules in early versus late spermatocytes in wild-type and *Ythdc2* cKO testes.

However, by P16, at which stage most germ cells had advanced to pachytene and most tubules had little immunofluorescence signal from anti-YTHDC2 in the *Ythdc2* cKO (Fig. 1O and 2Q), many genes were differentially expressed in *Ythdc2* cKO testes compared to controls (Fig. 5B and B′ and Fig. S5A and C). Further examination of the significantly differentially expressed genes in P16 *Ythdc2* cKO testes compared to *Cre* controls (437 genes total, adjusted p-value < 0.05) revealed that 313 genes were detected at lower levels in *Ythdc2* cKO testes (Table S1). Consistent with immunofluorescence experiments revealing proper protein expression of the meiotic markers SYCP3, SYCP1, γH2AX, HORMAD1, H1t and RAD51 in P16 *Ythdc2* cKO spermatocytes, expression of most meiotic genes were not significantly affected in P16 *Ythdc2* cKO mice compared to controls. However, a small subset of meiotic genes, including *Stag3, Terb1, Sun1, Psmc3ip, Syce1, Msh5, Zcwpw1* and *Mei1* were significantly under-expressed in *Ythdc2* cKO testes compared to controls (Table S1). Reciprocally, 124 genes were detected at higher levels (adjusted p-value < 0.05) in P16 *Ythdc2* cKO testes compared to *Cre* controls (Fig. 5B and B′). One of the most elevated genes in *Ythdc2* cKO testes compared to either control was *Meioc*.

Many of the genes significantly differentially expressed in P16 *Ythdc2* cKO testes compared to either the *Cre* or *flox/Δ* control had been identified as encoding RNAs directly bound by YTHDC2 in a recently published YTHDC2 CLIP-seq (UV cross-linking and immunoprecipitation followed by RNA sequencing) dataset from P15 testes (8) (Table S1), suggesting that at least some of the expression changes observed in the *Ythdc2* cKO testes may be a direct consequence of loss of function of *Ythdc2*. Both *Ythdc2* and *Meioc* RNAs were identified as directly bound by YTHDC2 by CLIP (8), suggesting that the YTHDC2-MEIOC protein-binding complex may help regulate its own levels of expression. Overall, 69% of the genes elevated in P16 *Ythdc2* cKO testes compared to *Cre* controls had been identified as encoding RNAs that were directly bound by YTHDC2 by CLIP, suggesting that YTHDC2 may act in mid to late spermatocytes to target the RNAs for degradation. In addition, 36% of the genes expressed at lower levels in P16 *Ythdc2* cKO testes than in *Cre* controls had also been identified as encoding RNAs directly bound by YTHDC2, including many of the highly down-regulated histone mRNAs such as *H2afx*, *H1f10* and *H4c12,* as well as genes required for sperm development, including *Prss50* (Fig. 5B′ and Fig. S5C), suggesting that YTHDC2 may be required to stabilize these RNAs.

A previous study showed by YTHDC2 CLIP in P15 testes that YTHDC2 binds to multiple sites at 5′ UTRs, coding sequences (CDS) and 3′ UTRs of ∼ 5,000 target genes, with a higher coverage at 3′ UTRs (8). Furthermore, the authors found that YTHDC2 bound several U-rich motifs in the 3′ UTR and suggested that YTHDC2 may be loaded onto the 3′ UTR of mRNAs using the U-rich motifs as multiple landing sites (8). Out of the list of genes encoding RNAs that were identified as possible direct YTHDC2-bound targets by CLIP (8), we observed that 867 genes were up-regulated (log2FC > 0.25) and 848 were down-regulated (log2FC < -0.25) in our *Ythdc2* cKO testes compared to *Cre* control testes (Fig 5C). Filtering this list of YTHDC2-bound target RNAs based on the enrichment for binding sites (BS) defined by the authors (8) to include only those with higher enrichment (BS ≥ 5) anywhere on the RNA, which would represent more frequent landing sites for YTHDC2, gave 388 up-regulated genes (log2FC > 0.25) and only 166 down-regulated genes (log2FC < -0.25) in *Ythdc2* cKO testes compared to *Cre* controls.

Because YTHDC2 was suggested to regulate RNA stability and/or degradation through binding the 3′ UTR of target RNAs, we examined the expression of genes encoding RNAs bound by YTHDC2 at the 3′ UTR. Indeed, from the 759 genes that showed YTHDC2 binding at the 3′ UTR by CLIP (BS ≥ 5), 193 were up-regulated (log2FC > 0.25) while only 60 were down-regulated (log2FC < -0.25) in *Ythdc2* cKO testes (Fig. 5D), suggesting that YTHDC2 binding to the 3′ UTR of many RNAs might target them for degradation. Li et al. (2022) found that in addition to 3′ UTRs, YTHDC2 can also bind CDS and 5′ UTRs, often in the same genes (8). To address whether binding of YTHDC2 at CDS or 5′ UTRs contributes to RNA degradation, we examined expression changes of genes encoding RNAs bound by YTHDC2 exclusively at the 3′ UTR, CDS or 5′ UTR (Fig. S6). We found 3.4x more genes up-regulated versus down-regulated in *Ythdc2* cKO testes when YTHDC2 bound only the 3′ UTR (BS ≥ 5) (Fig. S6C), while 1.6x more genes were up-regulated versus down-regulated when YTHDC2 bound only the CDS (BS ≥ 5) (Fig. S6B). Genes encoding transcripts bound only at the 5′ UTR appeared to have no specific enrichment for being up-regulated or down-regulated in *Ythdc2* cKO testes (Fig. S6A).

## DISCUSSION

The RNA-binding protein YTHDC2 is required for germ cell differentiation and survival at two different time points during meiotic progression. We and others have previously shown that an initial role of YTHDC2 in male germ cells entering meiotic prophase is to destabilize mitotic transcripts, including *Cyclin A2* (3, 5, 11). Failure to properly shut down the mitotic program upon meiotic entry caused germ cells in *Ythdc2* null mutant males to undergo an abnormal, mitosis-like division and apoptosis. Here we show, using a conditional knockout strategy to bypass the early functional requirement, that YTHDC2 has a second critical role in pachytene spermatocytes, where it is required for meiotic progression during the late pachytene stage. In testes subjected to conditional knockout of *Ythdc2* in early spermatocytes starting at P12 during the first wave of spermatogenesis, the germ cells did not express CYCLINA2 or undergo abnormal chromatin condensation, indicating that the cells had sufficient YTHDC2 function to successfully pass through the first requirement for YTHDC2 to prevent continued mitotic behavior, thus sparing the cells from early apoptosis. The germ cells in *Ythdc2* cKO males properly expressed and localized multiple meiotic markers and progressed to the late pachytene stage, unlike spermatocytes in *Ythdc2* null males, which died after leptotene. *Ythdc2* cKO spermatocytes reached a later stage than the stage IV mid-pachytene arrest often observed when defects occur in chromosomal synapsis or meiotic recombination (17). However, changes in expression or stability of certain meiotic RNAs were observed by P16, and the *Ythdc2* cKO spermatocytes failed to transition to the diplotene stage and began to undergo apoptosis by P18.

Because we induced *Ythdc2* knockout at P12 and P13 during the first wave of spermatogenesis and analyzed testes starting one day post the second tamoxifen injection, our experimental setup did not have later germ cell stages, allowing us to clearly observe the initial defects that occur upon loss of *Ythdc2* as male germ cells proceed synchronously through meiotic prophase. Our finding that *Ythdc2* cKO germ cells reached the late pachytene stage prior to undergoing apoptosis agreed with observations made by Liu and colleagues, who analyzed the effects of conditional knockout of *Ythdc2* in adult testes (6). However, because adult testes have germ cells at various stages of development, Liu et al. observed a mixture of both the early mitotic-to-meiotic phenotype as well as the later phenotype during late pachytene. As Liu et al. administered tamoxifen to adult mice for five consecutive days and analyzed the mice starting at two days post the final injection, some tubules in the adult *Ythdc2* cKO testes contained round and elongating spermatids, which had likely progressed past the pachytene stage during the spermatogenic waves before tamoxifen introduction (6). Liu et al. reported that nearly half of mid-pachytene spermatocytes and most late pachytene spermatocytes exhibited telomere clustering in their *Ythdc2* cKO model (6). We did not observe telomere clustering in *Ythdc2* cKO mid-pachytene spermatocytes at either P16 (*Ythdc2* cKO 0/271; control 0/216) or P18 (*Ythdc2* cKO 0/95; control 0/227). We did observe telomere clustering in *Ythdc2* cKO late pachytene spermatocytes, but in a much lower percentage of late pachytene cells in both P16 (*Ythdc2* cKO 17/142, 12%; control 0/41) and P18 (*Ythdc2* cKO 11/135, 8%; control 0/93) testes compared to Liu et al. (2021). This difference could be due to experimental timing (P16, P18 first wave versus adult), duration of tamoxifen exposure, genetic background, or the *Cre* driver used to knock out *Ythdc2* (*UBC-Cre^ERT2^* versus *Ddx4-Cre^ERT2^*).

By analyzing the effects of conditional knockout of YTHDC2 during the first wave of spermatogenesis, we were able to focus on the initial, likely more direct effects of loss of YTHDC2 function in spermatocytes. Our RNA-seq data from P16 *Ythdc2* cKO testes identified gene expression changes occurring soon after depletion of *Ythdc2* in spermatocytes. Many of the genes differentially expressed in *Ythdc2* cKO testes were previously identified as encoding RNAs that were directly bound by YTHDC2 based on CLIP (8, 9). Li et al. (2022) previously showed that YTHDC2 binding is enriched at the 3′ UTR of target RNAs (8). The authors also found that YTHDC2 recognizes U-rich motifs and quantified the number of binding sites in the target 3′ UTRs. Our analysis revealed that the presence of more YTHDC2 binding sites in the 3′ UTR correlated with an increase in expression of YTHDC2 target genes in P16 *Ythdc2* cKO testes compared to control, suggesting that binding of YTHDC2 at 3′ UTRs may facilitate degradation of these mRNAs in mid-to late-stage spermatocytes. As suggested by Li et al. (2022), having multiple YTHDC2 binding sites in the 3′ UTRs could increase the chance of overlap with binding sites for proteins that stabilize the transcripts, leading to YTHDC2 competing with and displacing the stabilizers from their targets.

While post-transcriptional regulation through action of YTHDC2 may alter RNA abundance for some RNAs bound by YTHDC2, we found that the changes in gene expression levels by RNA-seq in both the *Ythdc2* null mutant (3) and *Ythdc2* cKO were often quantitatively small, and many putative direct YTHDC2 targets identified by CLIP in both early (9) and late (8) spermatocytes showed no change in transcript levels at the time points we assessed. It is possible that binding of YTHDC2 could affect other processes, such as promoting or repressing translation or regulating localization or storage of RNAs that it binds.

YTHDC2 contains several RNA-binding domains, including an R3H domain, an OB fold and an RNA helicase core module (DEXDc and HELICc motifs). YTHDC2 has also been shown to have ATP-dependent, 3′ → 5′ RNA unwinding activity (5, 7). The RNA helicase domain appears to be required for function of YTHDC2 in late pachytene spermatocytes but not in early spermatocytes, as a point mutation in the YTHDC2 ATPase motif led to a late spermatocyte phenotype similar to the phenotype we and Liu et al. (2021) observed in *Ythdc2* cKO mice, rather than the earlier, mitosis-to-meiosis phenotype seen in *Ythdc2* null mutants (8, 9). This stage-specific effect of mutations that disable the RNA unwinding activity of YTHDC2 raises the possibility that one role of YTHDC2 in late pachytene spermatocytes could be to regulate translation. Previous studies suggested that YTHDC2 could promote translation of RNA targets, as YTHDC2 interacts with the small 40S ribosomal subunit (14). Binding of YTHDC2 to transcript coding sequences may help resolve mRNA secondary structure and promote translation *in vitro* (18). Ribosome profiling from P8 and P10 testes (9) indicated that YTHDC2 likely does not substantially regulate translation of its targets at the mitotic to meiotic switch.

However, it remains to be determined whether YTHDC2 acts through translational regulation in late spermatocytes. CLIP of YTHDC2 from P8, P10 and adult testes suggested that the RNAs bound by YTHDC2 vary across germ cell stages (9). Therefore, YTHDC2 action on target RNAs may differ depending on factors such as where on the RNA YTHDC2 binds (8, 9, 18), the number of YTHDC2 binding sites (Fig. 5D) or what other YTHDC2 protein interacting partners are present when the target RNA is bound.

Several studies have previously shown that YTHDC2 is part of a protein complex containing MEIOC (3, 10, 11), the RNA-binding protein RMB46 (8, 12), as well as the exoribonuclease XRN1 (7, 8, 14). To elucidate the mechanism(s) of action of YTHDC2 during early and late meiotic prophase, we immunoprecipitated YTHDC2 from P12 and P18 testis extracts followed by mass spectrometry to identify additional YTHDC2 interacting partners. Co-immunoprecipitation followed by western blot allowed us to distinguish RNA-independent and RNA-dependent YTHDC2 interacting partners in early and late spermatocytes. All the YTHDC2 interacting proteins we detected have known roles in regulating RNA biology. Several of the YTHDC2 binding partners were expressed at high levels in early spermatocytes, while others were more highly expressed in late spermatocytes, suggesting that YTHDC2 may associate with specific protein complexes at different cell stages, leading to diverse, stage-specific molecular outcomes on RNA targets. We recovered a small number of RBM46 peptides in our YTHDC2 IP-mass spec from P18 testes; however, RBM46 was not in the top 50 proteins in any of the replicates (Fig S4B). Similar RBM46 IP-mass spec from P12 and P21 mouse testes revealed a number of RBM46-associated proteins, several of which were also detected in the YTHDC2 protein complexes we identified, including MEIOC, PABPC1, MOV10 and UPF1 (12).

In our previous study examining the subcellular localization of YTHDC2, we showed that while YTHDC2 is distributed throughout the cytoplasm, it also concentrates in perinuclear puncta that stain for the P-body marker DCP1A (3). Here we show that, similar to YTHDC2, MEIOC protein is present throughout the cytoplasm as well as in perinuclear puncta that co-stain with the granule markers DCP1A and MIWI. Interestingly, *Meioc* RNA was expressed at higher levels in P16 *Ythdc2* cKO testes compared to either *Cre* control or *flox/Δ* control testes. In addition, localization of MEIOC protein to cytoplasmic puncta was strongly enriched in *Ythdc2* cKO pachytene spermatocytes compared to controls (Fig. 5E). One possibility is that YTHDC2 acts directly to move MEIOC protein out of granules in pachytene spermatocytes. Alternatively, other MEIOC binding partners could facilitate increased localization of MEIOC to granules in the absence of YTHDC2. Additional YTHDC2 protein binding partners identified in this study have also been shown to be components of P-bodies or other types of RNP granules, suggesting that YTHDC2 collaborates with RNA granule components. YTHDC2 could recruit target RNAs to granules where the RNAs are then degraded, translationally silenced, or stored, depending on the components of the granule. Alternatively, or in addition, YTHDC2 could return target RNAs from granules to the cytoplasm to be translated.

The conserved mouse proteins YTHDC2, MEIOC and RBM46 function together to regulate proper entry into and execution of the mitosis-to-meiosis transition. All three proteins play a critical role at the switch, as germ cells null mutant for any of the three proteins enter meiotic prophase but fail to shut off the mitotic program. As our current data reveal that YTHDC2 has an additional role in late spermatocytes, it would be interesting to examine whether MEIOC and RBM46 also function during late meiotic prophase, or if YTHDC2 may instead work with other protein partners to facilitate the transition from late pachytene to diplotene. Our carefully developed protocol for bypassing the early requirement for YTHDC2 function yet drastically knocking down its expression by the pachytene spermatocyte stage will facilitate future studies investigating YTHDC2, MEIOC and RBM46 and their roles as male germ cells progress through meiotic prophase.

## MATERIALS AND METHODS

### Mice

*Ythdc2* conditional knockout mice were generated according to the scheme depicted in Fig. S1. *Ythdc2^tp/tp^* mice (3) were crossed to C57BL/6N-Tg(CAG-Flpo)1Afst/Mmucd (RRID:MMRRC_036512-UCD) mice which removed the gene-trap cassette by Flp recombinase and reverted the mutation to wild type, with LoxP sites flanking exons 6 and 7 (*Ythdc2^flox^*). Removal of the gene-trap cassette was confirmed by PCR using the following primers: *Ythdc2* forward 5′arm: 5′-CTGAACATGTCTTATCCACAGTGC-3′, *Ythdc2* reverse 3′arm: 5′-CATCATCAAGAAGGTTACAACAGGC-3′, Neo forward: 5′-CAGCGCATCGCCTTCTATCGCC-3′. The *Ythdc2^flox^* mice were then crossed to mice carrying the *Ythdc2* null allele (*Ythdc2^+/Δ^*) (3) to generate *Ythdc2^flox/Δ^* mice. *Ythdc2^flox/Δ^* mice were crossed to either the inducible Cre strain *B6.Cg-Ndor1^Tg(UBC-Cre/ERT2)1Ejb^/1J* (RRID:IMSR_JAX:007001) (15) or to *Tg(Spo11-Cre)1Rsw/PecoJ* (RRID:IMSR_JAX:032646) (16). Mice were genotyped by PCR using the following primers: *Cre* forward: 5′-TGGGCGGCATGGTGCAAGTT-3′, *Cre* reverse: 5′-CGGTGCTAACCAGCGTTTTC-3′, *Ythdc2* forward: 5′-CGAGTGCTGCCTTGGATGTGAACC-3′, *Ythdc2* reverse: 5′-GGATTTTGACAGCCTTGAGCCTGGG-3′, targeting cassette (*LacZ*) forward: 5′-GAATTATGGCCCACACCAGTGGCG-3′. All experiments were approved by the Stanford University Animal Care and Use Committee and performed according to NIH guidelines.

### *Ythdc2* knock out with Tamoxifen

Tamoxifen (10 mg/ml, Cayman Chemical) was dissolved in corn oil by incubating at 55°C for 40 min with constant rotation and then 0.1 mg/g body weight was injected intraperitoneally using a 27G needle into *UBC-Cre^ERT2^; Ythdc2^flox/Δ^*(*Ythdc2* cKO), +/+; *Ythdc2^flox/Δ^* (*flox/Δ* control) or *UBC-Cre^ERT2^; Ythdc2^flox/+^* (*Cre* control) male mice at P12 and P13 at the same time each day.

### Histology

Mouse testes were fixed overnight at room temperature (RT) in bouins fixative and embedded in paraffin. Paraffin-embedded samples were cut at 5-6 μm thickness and the sections were stained with periodic acid-Schiff (PAS).

### Immunofluorescence

For immunofluorescence staining, mouse testes were fixed with 4% paraformaldehyde overnight at 4°C, embedded in paraffin and cut to 5-6 μm thickness. After rehydrating in an ethanol gradient, heat-mediated antigen retrieval was performed on the sections using sodium citrate buffer (10 mM sodium citrate, 0.05% Tween-20, pH 6.0). Sections were then permeabilized with phosphate-buffered saline + 0.1% TritonX-100 (PBST) for 45 min at RT, followed by 1 hr incubation with blocking buffer (10% BSA in PBST) at RT. Sections were incubated with primary antibody overnight at 4°C and then incubated with Alexa Fluor-conjugated donkey secondary antibodies (1:400, Molecular Probes) at RT for 2 hrs and then mounted in VECTASHIELD medium with 4′,6-Diamidino-2-phenylindole (DAPI) (Vector Lab Inc.).

When immunolabeling with antibodies from the same host species, following incubation with primary antibody overnight at 4°C, slides were washed 3 times in PBS, and then incubated with HRP anti-rabbit IgG (1:500 in 10% BSA in PBST) for 1 hr at RT. Following incubation, TSA plus cyanine 3 solution (dilute TSA-Cy3 1:250 in 100mM Boric Acid, pH 8.5 containing 0.003% H_2_O_2_) (Akoya Biosciences) was added to the tissue sections and incubated for 10 min at RT. Slides were then washed twice in PBS and antigen retrieval using sodium citrate buffer was performed a second time. Sections were blocked and then incubated with primary and secondary antibodies as described above.

### Antibodies

Rabbit anti-YTHDC2 (1:500 or 1:2,000 when TSA used, A303-026A, (RRID:AB_10754785), Bethyl laboratories), mouse anti-SYCP3 (1:200, clone [Cor 10G11/7], (RRID:AB_10678841), Abcam), rabbit anti-SYCP3 (1:500, NB300-232, (RRID:AB_2087193), Novus Biologicals), rabbit anti-SYCP1 (1:500, NB300-229SS, (RRID:AB_922549), Novus Biologicals), rabbit anti-HORMAD1 (1:1,000, 13917-1-AP, (RRID:AB_2120844), Proteintech), guinea pig anti-H1t (1:500, Mary Ann Handel (19)), human anti-centromere (1:50, 15-234, (RRID:AB_2939058), Antibodies Incorporated), rabbit anti-RAD51 (1:200 [N1C2], (RRID:AB_1951602) , GeneTex), goat anti-VASA (1:200, AF2030, (RRID:AB_2277369), R&D Systems), mouse anti-phospho-Histone H2A.X (Ser139) (1:500 clone [JBW301], (RRID:AB_309864), Millipore), rabbit anti-CYCLIN A2 (1:250 clone [EPR17351], (RRID:AB_2890136), Abcam), rabbit anti-MEIOC (1:1,000 IF, David Page (11)), mouse anti-DCP1A (1:250 clone [3G4], (RRID:AB_1843673), Sigma), goat anti-PIWIL1 (1:200, AF6548, (RRID:AB_10971944), Novus Biologicals). Terminal deoxynucleotidyl transferase dUTP nick end labeling (TUNEL) staining was done following manufacturer’s instruction (In Situ cell Death Detection Kit TMR red, Roche).

The following antibodies were used for immunoblot: rabbit anti-YTHDC2 (1:1,000, A303-026A, (RRID:AB_10754785), Bethyl laboratories), rabbit anti-MEIOC (1:2000 immunoblot, David Page), goat anti-RENT1/UPF1 (1:2,000, A300-038A, (RRID:AB_2288326), Bethyl laboratories), rabbit anti-EWS (1:2,000, A300-417A, (RRID:AB_420957), Bethyl laboratories), rabbit anti-MOV10 (1:2,000, A301-571A, (RRID:AB_1040002), Bethyl laboratories), rabbit anti-G3BP2 (1:2,000, A302-040A, (RRID:AB_1576545), Bethyl laboratories), rabbit anti-YBX1 (1:2,000, A303-231A, (RRID:AB_10951283), Bethyl laboratories), mouse anti-PTBP1 (1:2,000, Thermo), rabbit anti-PABPC1 (1:1,000, ab21060, (RRID:AB_777008, Abcam), rabbit anti-FMR1 (1:2,000, A305-200A, (RRID:AB_2631593), Bethyl laboratories), rabbit anti-CAPRIN1 (1:2,000, HPA018126, (RRID:AB_1849929), Sigma).

YTHDC2 antibodies A303-026A (used for immunofluorescence and immunoblot) and A303-025A (used for immunoprecipitation) were previously validated for YTHDC2 specificity by testing the antibodies side-by-side on wild-type and *Ythdc2* null mutant testes (3).

### Germ cell spreads

Chromosome spreads from P16 and P18 testes were performed based on a previously published protocol (20). Briefly, testis tubules were incubated in a hypotonic extraction buffer pH 8.2 (30 mM Tris-HCL pH 8.0, 50 mM sucrose, 17 mM trisodium citrate dehydrate, 5 mM EDTA, 0.5 mM DTT, 0.5 mM PMSF) for 30 min. Testis tubules were then broken apart in 100 mM sucrose pH 8.2 and then spread on a slide dipped in fixation buffer pH 9.2 (1% PFA and 0.15% Triton X-100). Slides were incubated overnight in a humidified chamber at RT and air-dried. Slides were washed twice in 0.4% Photoflo (Kodak) and air-dried.

### Telomere clustering quantification

Germ cell spreads from P16 and P18 *Ythdc2* cKO and *flox/Δ* control testes were stained with anti-SYCP3 and anti-H1t to distinguish early, mid, and late pachytene substages. Germ cells were also stained with anti-CREST to mark the centromere. The centromere is positioned at one end of the chromosome adjacent to the telomere in mouse, allowing us to quantify the number of mid and late pachytene spermatocytes that showed clustering. N=2 biological replicates performed for each time point and genotype.

### Co-immunoprecipitation from testes

Protein A Dynabeads (50 μL/IP, Invitrogen) were blocked briefly in 3% BSA in PBS + 0.1% tween 20 (PBST) and then incubated at 4°C overnight in lysis buffer (20 mM Tris-HCL, 135mM NaCl, 10% glycerol, 1% Nonidet P-40, 5 mM EDTA) plus 2 μg YTHDC2 antibody (A303-025A, Bethyl laboratories). Beads were washed three times with 0.2 M triethanolamine, pH 8.2 for 5 min at RT, then incubated with dimethylprimelimidate (DMP, 5.4 mg DMP per mL 0.2 M triethanolamine, Sigma) for 30 min at RT to crosslink the antibody to the beads. Beads were then washed once with 50 mM Tris for 15 min, followed by three 5 min washes with PBST. Crosslinked beads can be stored overnight at 4°C or used immediately in the next step. To remove un-crosslinked antibody, beads were quickly washed twice with 100mM glycine, pH 2.5, then blocked in 3% BSA PBST for 30 min followed by three quick PBST washes.

Testes were dissected from P12 and P18 wild-type mice. Testis extracts were prepared by mechanically disrupting testes in lysis buffer (20 mM Tris-HCL, 135 mM NaCl, 10% glycerol, 1% Nonidet P-40, 5 mM EDTA, 1 mM PMSF, 1x Complete protease inhibitor, and either (A) 100 U/mL RNAse-OUT or (B) 50 µg/mL RNAse A). Extracts were incubated in lysis buffer for 20 min at 4°C, spun for 20 min, and then precleared with uncoupled, 3% BSA-blocked Protein A Dynabeads for 1 hr at 4°C. Testis extracts were then added to the crosslinked YTHDC2 antibody - Protein A Dynabeads and incubated for 3 hrs at 4°C while rotating. Beads were washed three times in lysis buffer for 5 min each wash, and incubated at 70°C for 30 min in elution buffer (1% SDS, 10 mM EDTA, 50 mM Tris pH 8.0, 1 mM PMSF, 1x Complete protease inhibitor) with frequent mixing. Samples were resolved on a 10% SDS-PAGE gel (Bio-Rad) and transferred onto polyvinylidene fluoride (PVDF) membranes. Blots were incubated in primary antibodies overnight at 4°C and then IgG HRP secondary antibodies (1:10,000) for 2 hrs at RT. Blots were developed using western lightning ECL detection reagent.

### Immunoprecipitation Mass Spectrometry

Testes were dissected from P12 and P18 wild-type and *Ythdc2* mutant mice. YTHDC2 or IgG immunoprecipitations were performed as described above with slight modifications: the uncoupled, preclear Protein A Dynabeads were washed with PBST to remove unbound BSA prior to adding to the testis lysate, and the crosslinked YTHDC2 or IgG antibody - Protein A Dynabeads were not blocked with 3% BSA following DMP crosslinking. Testis extracts were precleared with uncoupled beads for 3 hrs at 4°C followed by incubation with the crosslinked YTHDC2 antibody - Protein A Dynabeads for 3 hrs at 4°C with continuous rotation. Beads were washed three times in lysis buffer for 10 min each wash, and then incubated in elution buffer as previously described above. Samples were resolved on a 10% SDS-PAGE gel (Bio-Rad). The samples were allowed to run about 1 cm into the gel and the gel was then fixed for 1 hr in fixing solution (45:45:10 / water:methanol:acetic acid) with gentle agitation. The samples were cut out of the gel and stored at 4°C until being processed at the Stanford University Mass Spectrometry Facility. YTHDC2 IPs in P12 *Ythdc2* null mutant mice and IgG IPs in P12 and P18 wild-type mice served as negative controls. N=3 biological replicates were performed.

### RNA-seq from testes

Testes were collected from P14 and P16 *UBC-Cre^ERT2^; Ythdc2^flox/Δ^* (*Ythdc2* cKO ) mice as well as both +/+; *Ythdc2^flox/Δ^*(*flox/Δ* control) and *UBC-Cre^ERT2^; Ythdc2^flox/+^*(*Cre* control) mice following tamoxifen injections at P12 and P13. One testis was snap frozen and the other was fixed in 4% formaldehyde to confirm *Ythdc2* knockout. Total RNA was isolated from frozen testes using the RNeasy Plus Mini Kit (Qiagen), according to the manufacturer’s instructions. Ribosomal RNAs were depleted from 1 µg of total RNA using either TruSeq Stranded Total RNA kit Ribo-Zero components (Illumina) (*flox/Δ* controls and *Ythdc2* cKO replicate 1&2) or the NEBNext rRNA Depletion Kit v2 (*Cre* controls and *Ythdc2* cKO replicate 3). Ribosomal-depleted RNA was cleaned up using Agencourt RNAClean XP Beads (Beckman Coulter) and efficiency of rRNA depletion was confirmed by bioanalyzer. Library preparation for high-throughput sequencing was performed on 100 ng of rRNA depleted RNA using the NEBNext Ultra II Directional RNA Library Prep Kit for Illumina (New England Biolabs). Sequencing was done with a NextSeq 500/550 High Output Kit v2.5. Each condition had n=2-4 biological replicates.

### RNA expression analysis

Libraries were separated by barcode and adapters and low-quality bases were trimmed using trimGalore (21). Reads were mapped to the mouse genome (GRCm39/mm39) using STAR (RRID:SCR_004463) (22). Reads for each transcript were extracted using HTSeq (RRID:SCR_005514) (23). Differential gene expression was calculated using DESeq2 (RRID:SCR_015687) (24). Batch effects were accounted for in the design of the experiment in DESeq2 when comparing samples that used different methods of ribosomal depletion. Batch effects were removed using Limma to generate the PCA plot (Fig. S5A). YTHDC2-bound RNAs identified by YTHDC2 CLIP-seq from P15 testes by Li et al. (2022) (GEO accession GSE196427) (8) were compared to gene expression changes observed in P16 *Ythdc2* cKO testes from this study. YTHDC2 binding sites (BS) on the 5′ UTRs, CDS and 3′ UTRs of target RNAs were identified by Li et al. (2022) (8).

### Statistical analysis

For phenotypic analysis, sample sizes (n) for each experiment are given in the respective figure legends. Sample sizes for telomere clustering quantification, RNA-seq, and Mass Spec experiments, are provided in the methods. Sample sizes used were similar to what is generally utilized in the field. Each animal was considered a biological replicate. All bar graphs are presented as means ± SEM. Statistical differences between two groups were analyzed using unpaired two-tailed t tests; p<0.05 was considered statistically significant.

### Data availability

RNA-seq data have been deposited in the Gene Expression Omnibus (GEO) under accession number GSE222283.

## ACKNOWLEDGEMENTS

We thank members of the Fuller laboratory for many helpful discussions, especially Neuza Matias for help with RNA-seq and CLIP-seq analysis, Sarah Stern for help with data analysis and presentation, and Catherine Baker for helpful feedback throughout the project. We would also like to thank David Page for providing the MEIOC antibody, Mary Handel for providing the H1t antibody and Georgi Marinov for running the high-throughput sequencing. We also thank the Stanford Comparative Medicine Animal Histology Services for preparing histological sections, the Stanford Protein and Nucleic Acid Facility for RNA quality control analysis, and the Vincent Coates Foundation Mass Spectrometry Laboratory, Stanford University Mass Spectrometry core.

