## Supplemental Figures for "YTHDC2 serves a distinct late role in spermatocytes during germ cell differentiation"

**SUPPORTING INFORMATION**

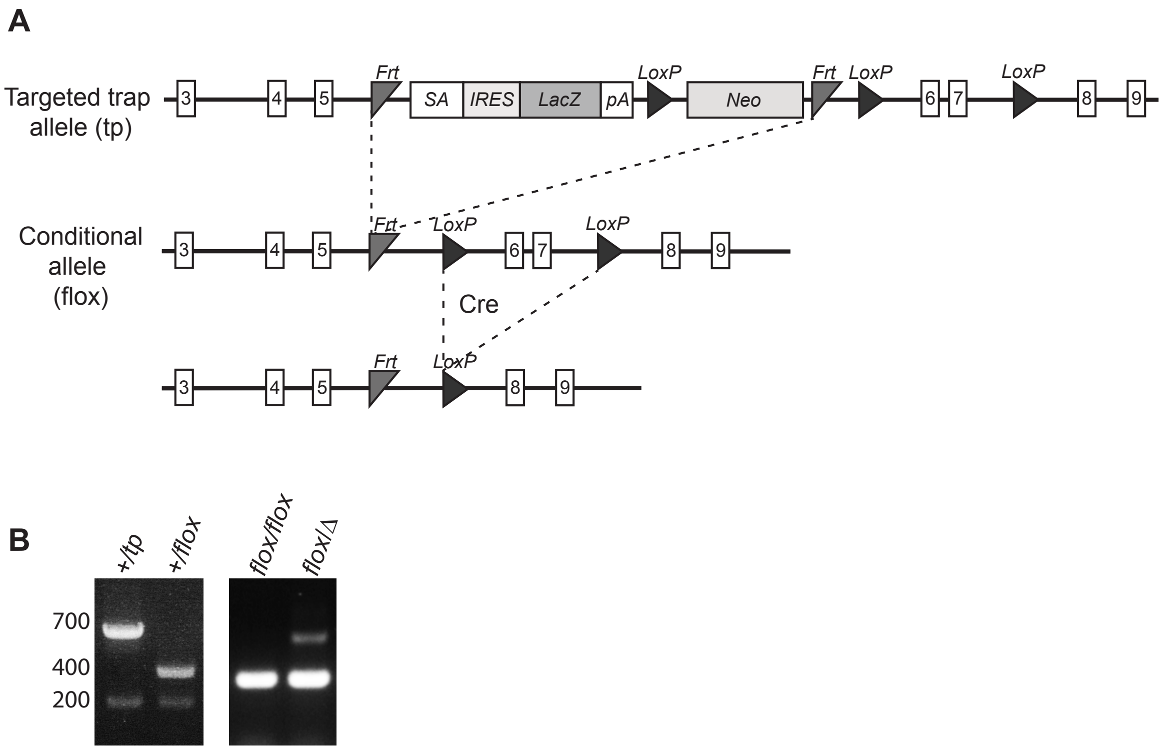


**Fig. S1. Generation of the *Ythdc2* conditional knockout mouse.** (A) Diagram of the generation of the *Ythdc2* conditional allele (*flox*) from the targeted trap allele (*tp*) through excision of the LacZ reporter and the Neomycin cassette. Cre-mediated deletion of exons 6 and 7 occurs in the *Ythdc2* floxed mice upon exposure to Cre recombinase. SA, splice acceptor; IRES, internal ribosome entry site; LacZ, β-galactosidase gene; pA, polyadenylation sequence; LoxP, Cre recombinase recognition site; FRT, flippase recognition site. (B) Mouse genotyping by PCR analysis of DNA purified from mice heterozygous for the *Ythdc2* targeted trap allele (*+/tp*), heterozygous for the conditional allele (*+/flox*), homozygous for the conditional allele (*flox/flox*) and heterozygous for the conditional and null allele (*flox/*Δ).

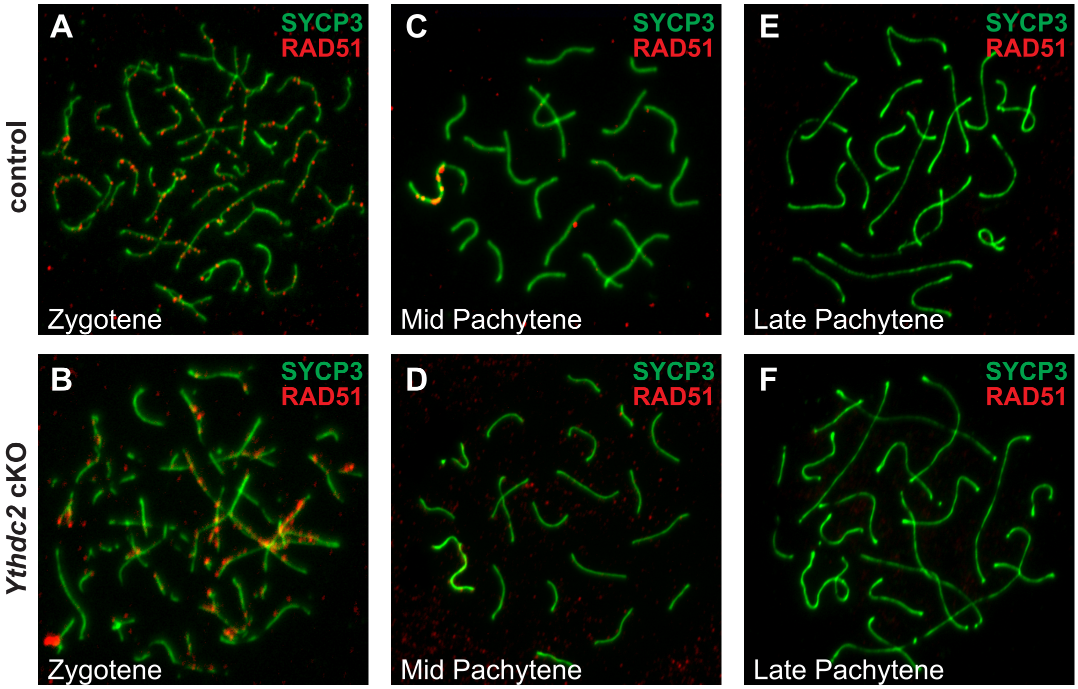


**Fig. S2. Normal expression pattern of RAD51 in early and late spermatocytes.** (A-F) Immunofluorescence images of germ cell spreads from P18 tamoxifen-treated control (*+/+; Ythdc2*$^{flox/\Delta}$, top) and *Ythdc2* cKO (*UBC-CreERT2; Ythdc2*$^{flox/\Delta}$, bottom) mice stained with anti-SYCP3 (green) and anti-RAD51 (red) antibodies.

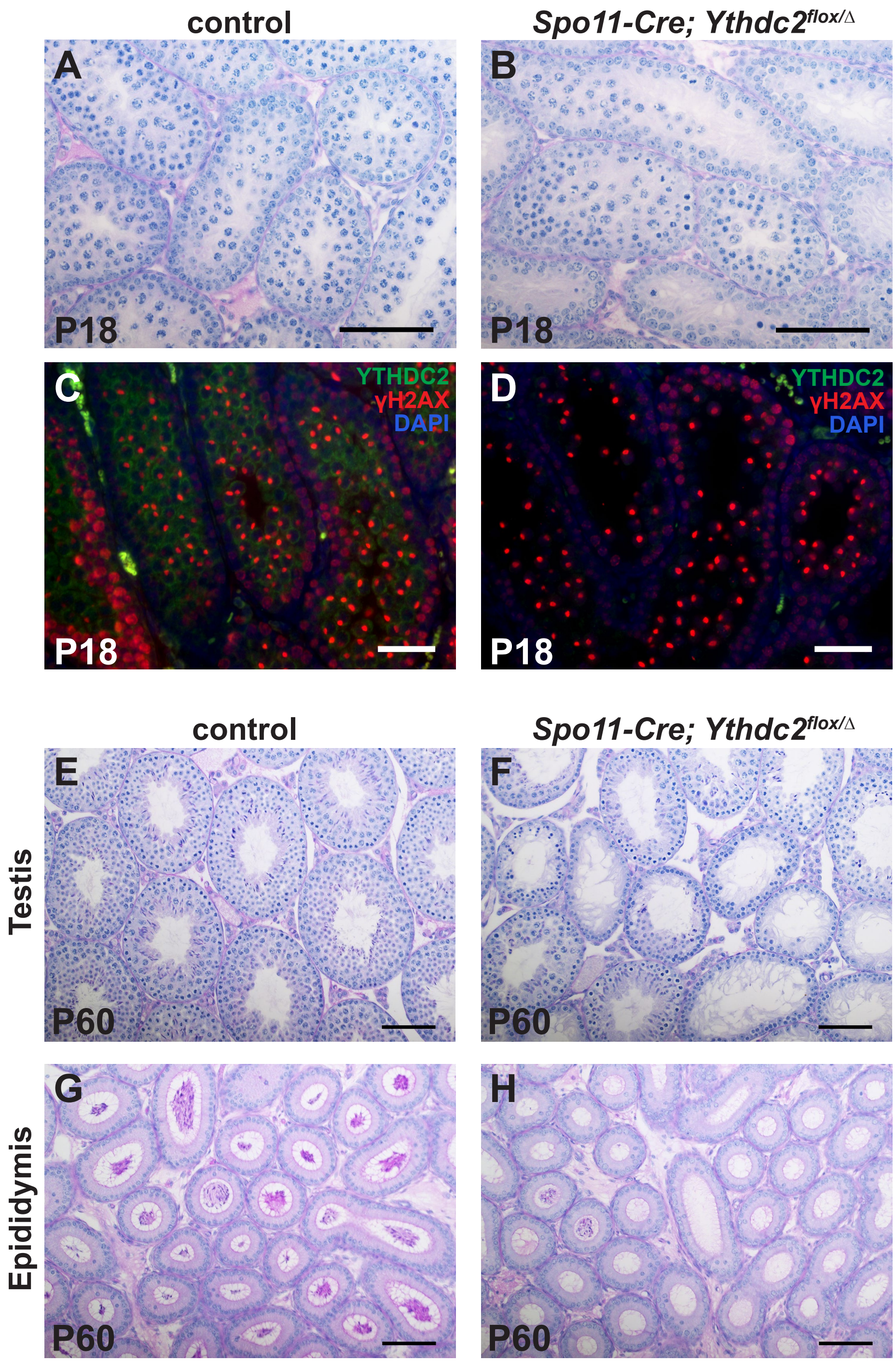


**Fig. S3. Knocking out *Ythdc2* using the *Spo11-Cre* driver leads to a similar phenotype as the inducible driver strategy.** (A and B) Cross-sections of mouse seminiferous tubules stained with periodic acid-Schiff (PAS) from P18 (A) control (*Spo11-Cre; Ythdc2*$^{flox/+}$) and (B) *Spo11-Cre; Ythdc2*$^{flox/\Delta}$ mice. (C and D) Immunofluorescence images of P18 (C) control and (D) *Spo11-Cre; Ythdc2*$^{flox/\Delta}$ mouse testis tubule cross-sections stained with anti-YTHDC2 (green) and anti-γH2AX (red) antibodies as well as DAPI to mark the DNA (blue). (E and F) Cross-sections of mouse seminiferous tubules stained with PAS from P60 (E) control and (F) *Spo11-Cre; Ythdc2*$^{flox/\Delta}$ mice. (G and H) Cross-sections of mouse epididymis stained with PAS from P60 (G) control and (H) *Spo11-Cre; Ythdc2*$^{flox/\Delta}$ mice. Scale bars: 50μm.

**A** Proteins detected in P12 YTHDC2 complexes

| Protein | Number of spectra |
|---|---|
| PABPC1 | 20 |
| FMR1 | 9 |
| PTBP1 | 8 |
| UPF1 | 5 |
| G3BP2 | 5 |
| MOV10 | 4 |
| MEIOC | 3 |
| CAPRIN1 | 3 |
| EWS | 2 |

**B** Proteins detected in P18 YTHDC2 complexes

| Protein | Number of spectra |
|---|---|
| PABPC1 | 40 |
| MEIOC | 12 |
| UPF1 | 10 |
| YBX2 | 8 |
| PTBP1 | 6 |
| G3BP2 | 6 |
| FMR1 | 5 |
| CAPRIN1 | 5 |
| EWS | 5 |
| MOV10 | 3 |
| RBM46* | 2 |

RNA-independent

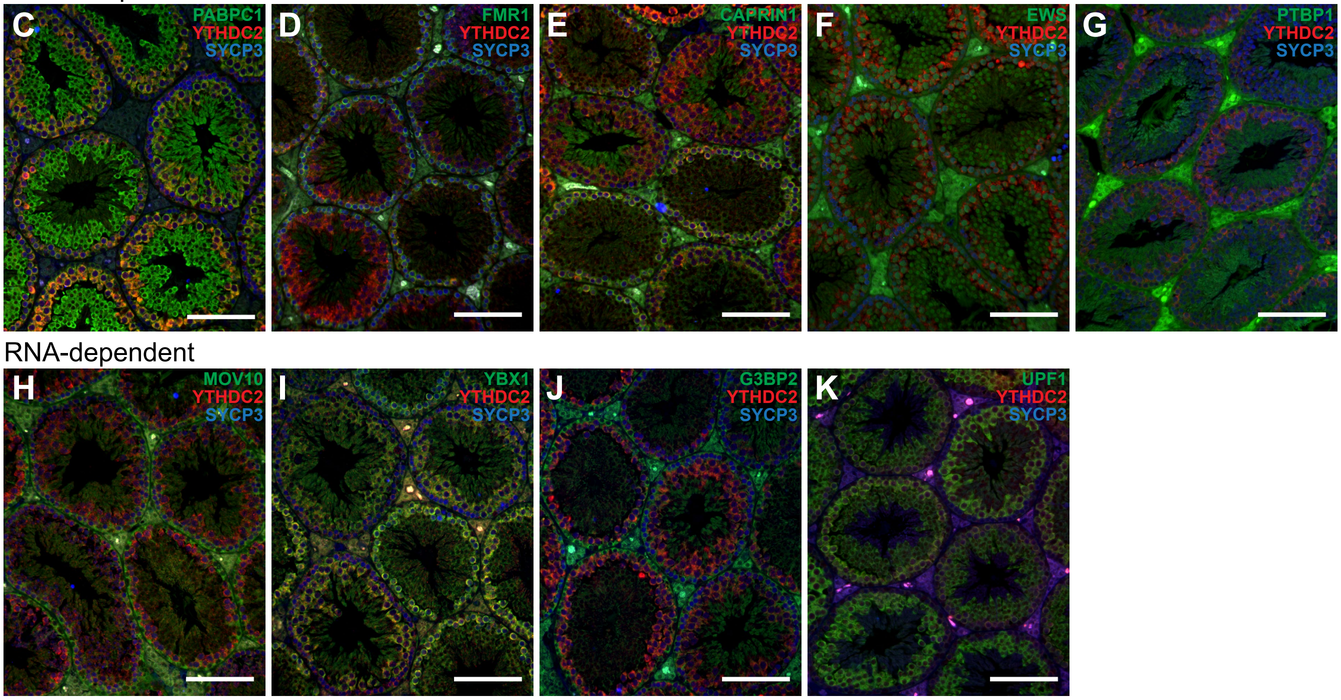


**L**

| Protein | Function | Protein expression pattern in spermatocytes by immunofluorescence |
|---|---|---|
| MEIOC | RNA stability | Leptotene, zygotene & pachytene (cytoplasmic) |
| PABPC1 | RNA stability, translation initiation & RNA degradation | Low in leptotene & zygotene / high in pachytene (cytoplasmic) |
| FMR1 | RNA stability, translation, RNA processing & RNA transport | Leptotene, zygotene & early pachytene (cytoplasmic) |
| CAPRIN1 | Translation & RNA transport | Pachytene (cytoplasmic) |
| EWS | Transcription co-regulator activity & RNA binding | Leptotene, zygotene (cytoplasmic, nuclear) & pachytene (nuclear) |
| PTBP1 | Alternative splicing, RNA localization, stability & translation | Leptotene & zygotene (cytoplasmic) |
| MOV10 | RNA stability, translation & RNA cleavage/gene silencing by miRNA | Leptotene & zygotene (cytoplasmic) |
| YBX1 | RNA stability, RNA processing, translational repression & splicing | Pachytene (cytoplasmic) |
| G3BP2 | Scaffold protein, granule assembly & RNA transport | Leptotene & zygotene (cytoplasmic) |
| UPF1 | Nonsense mediated mRNA decay & RNA destabilization | Pachytene (cytoplasmic) |

**Fig. S4. YTHDC2 protein candidate binding partners in spermatocytes.**
(A and B) Top YTHDC2-associated protein candidates identified by YTHDC2 IP followed by mass spec from (A) P12 and (B) P18 mouse testes. (* A small number of peptides were seen for the previously identified YTHDC2 binding partner RBM46 in P18 testes. However, RBM46 was not in the top 50 proteins in any replicate.) (C-K) Immunofluorescence images of adult mouse testis tubule cross-sections stained for YTHDC2 (red), SYCP3 (blue) and either (C) PABCP, (D) FMR1, (E) CAPRIN1, (F) EWS, (G) PTBP1, (H) MOV10, (I) YBX1, (J) G3BP2 or (K) UPF1 (green). Scale bars: 100μm. (L) Table listing the protein expression pattern determined by immunofluorescence staining and the function of the YTHDC2 binding partners.

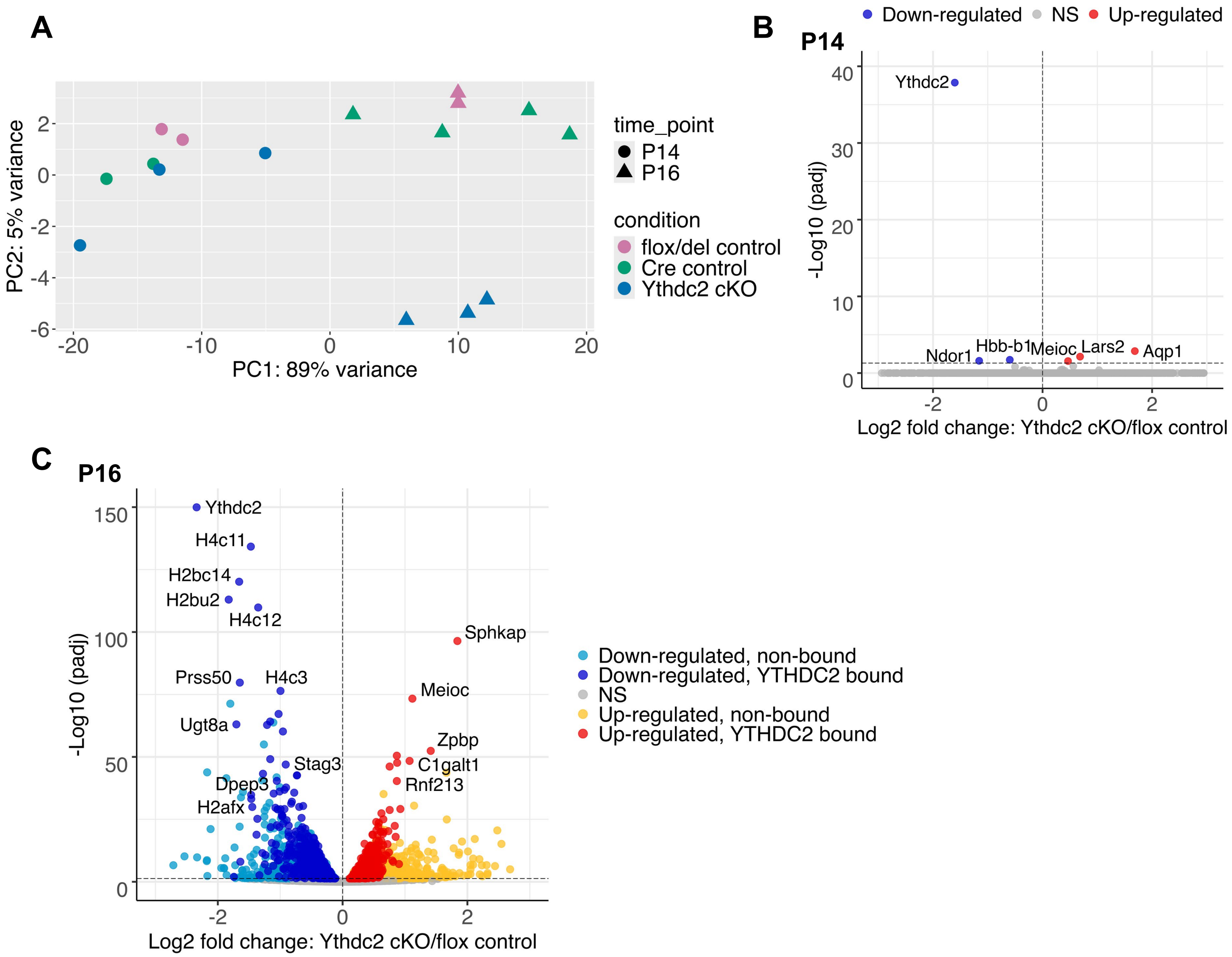


**Fig. S5. Early transcript changes following knockout of *Ythdc2* in spermatocytes.** (A) Principal component analysis (PCA) of P14 and P16 testes from *Ythdc2* cKO (*UBC-CreERT2; Ythdc2*$^{flox/\Delta}$), *flox/*Δ control (+/+*; Ythdc2*$^{flox/\Delta}$), and *Cre* control (*UBC-CreERT2; Ythdc2*$^{flox/+}$) samples. Principal component 1 (PC1) splits the samples by time point. Principal component 2 (PC2) splits the samples by genotype. (B and C) Volcano plots representing significantly differentially expressed genes (adjusted p-value < 0.05, horizontal dashed line) in *Ythdc2* cKO (*UBC-CreERT2; Ythdc2*$^{flox/\Delta}$) testes relative to *flox/*Δ control (+/+*; Ythdc2*$^{flox/\Delta}$) testes at (B) P14 and (C) P16. (B) (Red) genes up-regulated and (Blue) genes down-regulated in *Ythdc2* cKO testes relative to *flox/*Δ control testes. (C) Genes up and down-regulated in *Ythdc2* cKO testes color-coded based on whether the encoded transcripts were previously identified as bound by YTHDC2 based on published YTHDC2 CLIP data (1). (Light blue) down-regulated, non-bound; (Dark blue) down-regulated, YTHDC2 bound; (Yellow) up-regulated, non-bound; (Red) up-regulated, YTHDC2 bound.

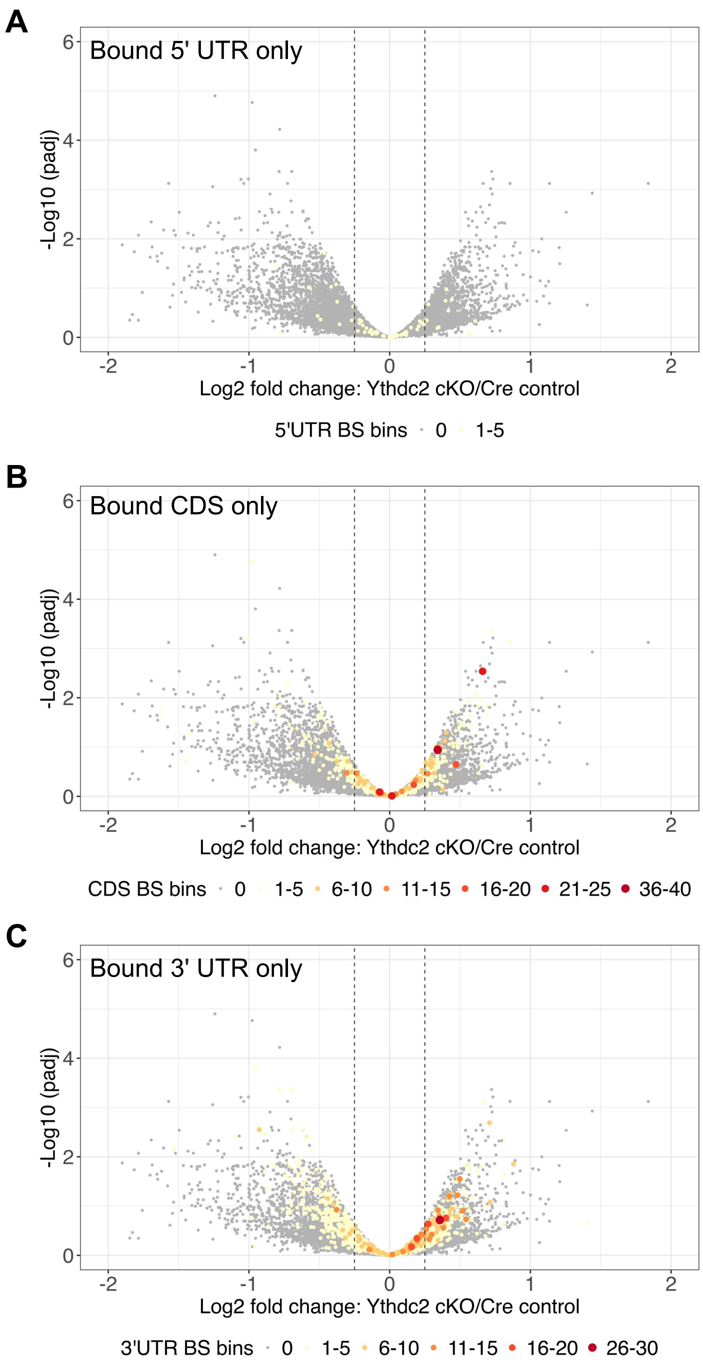


**Fig. S6. Expression changes for RNAs bound by YTHDC2 at the 5′ UTR only, CDS only or 3′ UTR only.** (A-C) Volcano plots depicting fold changes in gene expression in P16 *Ythdc2* cKO (*UBC-CreERT2; Ythdc2*$^{flox/\Delta}$) testes relative to *Cre* control (*UBC-CreERT2; Ythdc2*$^{flox/+}$) testis. Genes identified through CLIP as bound by YTHDC2 only at their (A) 5′ UTR, (B) coding sequence (CDS) or (C) 3′ UTR are highlighted using a gradient color scale representing the number of YTHDC2 binding sites (BS) as determined by Li et al. (2022) (1). Dashed gray lines indicate Log2FC -0.25 and Log2FC 0.25.
